# Subgenomic SARS-CoV-2 replicon and reporter replicon cell lines enable ultrahigh throughput antiviral screening and mechanistic studies with antivirals, viral mutations or host factors that affect COVID-19 replication

**DOI:** 10.1101/2021.12.29.474471

**Authors:** Shuiyun Lan, Philip R. Tedbury, Yee Tsuey Ong, Raven Shah, Ryan L. Slack, Maria E. Cilento, Huanchun Zhang, Haijuan Du, Nicole Lulkin, Uyen Le, Karen A. Kirby, Ivo Melcak, William A. Cantara, Emerson A. Boggs, Stefan G. Sarafianos

## Abstract

Replicon-based technologies were used to develop reagents and assays for advanced drug discovery efforts against severe acute respiratory syndrome coronavirus 2 (SARS-CoV-2), and for examining all facets of the SARS-CoV-2 replication cycle at reduced biocontainment level. Specifically: a) 21 replicons were cloned in bacterial artificial chromosomes (BACs) and delivered as transfectable plasmid DNA or transcribed RNA in various cell types. Replicons carrying mutations that affect the activity or antiviral susceptibility of SARS-CoV-2 enzymes were used to establish utility for mechanistic studies while reducing the community risks associated with gain-of-function studies in fully infectious virus. b) A BHK-21 stable cell line harboring SARS-CoV-2 replicon was generated and characterized in robust high/ultra-high throughput assays of antiviral efficacy with orthogonal SARS-CoV-2 replication reporter genes (Nano luciferase and enhanced green fluorescent protein-eGFP); the estimated antiviral potencies in the fully infectious SARS-CoV-2 system and in the transient or stable replicon systems were similar. HEK293 and Calu1 stable cell lines expressing SARS-CoV-2 replicon have also been prepared. Finally, c) we generated trans-encapsidated replicons by co-expression with SARS-CoV-2 structural proteins, thus producing single-round infectious SARS-CoV-2 virus-like particles that are able to transduce susceptible cell types and have expanded utility to enable study of virion assembly and entry into target cells. Hence, these SARS-CoV-2 replicon-based reagents include a novel approach to replicon-harboring cell line generation and are valuable tools that can be used at lower biosafety level (BSL2) for drug discovery efforts, characterization of SARS-CoV-2 and variant evolution in the COVID-19 pandemic, mechanisms of inhibition and resistance, and studies on the role of SARS-CoV-2 genes and host dependency factors.

## INTRODUCTION

As of the end of 2021, we are at the peak of the global COVID-19 pandemic, a respiratory disease with more than 280 million confirmed cases (record-high single-day infections on 12/23/2021 of 982,000 cases) and over 5.4 million fatalities around the world (1). The causative agent of COVID-19 (2, 3) is Severe Acute Respiratory Syndrome Coronavirus 2 (SARS-CoV-2), an enveloped, positive-sense, single-stranded RNA betacoronavirus of the order Nidovirales, family Coronaviridae. Prior to SARS-CoV-2, illness caused by endemic coronaviruses (4), and the outbreaks of the original SARS-CoV in 2002 (5-7) and Middle-East Respiratory Syndrome CoV (MERS-CoV) in 2012 (8) (9), highlight the threat coronaviruses pose to human health.

The vaccines currently in use for the prevention of SARS-CoV-2 infection (10, 11) have proven to be effective at reducing the hospitilization and mortality from COVID-19, and are likely to continue to be a major part of the response to this and future outbreaks. However, vaccination strategies face multiple challenges, including the limited global vaccine supply, the difficulty in achieving the theoretical herd-immunity threshold at current vaccination rates, the reported waning of immunity against the virus a few months after vaccination, and the emergence of SARS-CoV-2 variants that spread very efficiently and are extensively mutated at the Spike surface glycoprotein, which has been the main target for vaccine development (12, 13) (14-16). Therefore, effective antivirals that target viral proteins less likely to mutate will be an critical component of the global response to coronaviral outbreaks.

The nucleoside analogue remdesivir (RDV) was the first SARS-CoV-2-targeting antiviral approved by FDA for treatment of COVID-19 patients requiring hospitalization (reviewed in (17)). However, it can only be injected at a hospital setting and its efficacy has been questioned (18, 19). Recently, another nucleoside analog was approved: molnupiravir (EIDD-2801 or MK-4482), which is the orally available pro-drug of the β-d-N4-hydroxycytidine (20). Molnupiravir reduces hospitalization or death by 30% (21, 22). Viral proteases have also been excellent targets for the development of antiviral drugs (23, 24). Several have been reported to have activity *in vitro* or are in clinical trials (25-28), and one formulation, Paxlovid (nirmatrelvir with ritonavir) (29, 30), was recently approved for use in the USA (31). Paxlovid was reported to reduce the risk of hospitalization or death by 89% in non-hospitalized high-risk adults (clinical trial NCT04960202) (32). These major targets, and several other enzymatic activities are essential to replicon activity, allowing replicons to be used for initial screening for drug discovery, and characterization of mechanism of action and potential mechanisms drug of resistance, following mutation of the viral genome.

There is clearly a need for discovery and development of novel drugs that target SARS-CoV-2 replication. The virus is readily cultured and infectious clones have been produced with reporter viruses to facilitate such studies (33, 34). However, a challenge for such studies is the requirement of handling infectious SARS-CoV-2 in biosafety level-3 (BSL3) facilities, which limits the number of academic and pharmaceutical laboratories able to contribute to the search for effective therapeutics. Subgenomic replicon systems provide biologically safe models that recapitulate a large part of the viral replication cycle and can play a major role in the discovery and development of antivirals, in addition to providing an invaluable tool for basic research into viral replication (35-37).

The general strategy of producing subgenomic replicons involves removing the structural protein coding sequences and replacing them with reporters and/or selectable markers, while retaining all proteins and any nucleic acid sequences required for genome replication. The genomic RNA (gRNA) of coronaviruses is flanked by 5’ and 3’ untranslated regions (UTR); the first two thirds of the genome codes for the non-structural proteins (nsp) in the form of ORF1a and ORF1b polyproteins (reviewed in (38, 39)), the final third codes for the structural proteins (spike S, Envelope E, Membrane M, and Nucleocapsid N), as well as several putative ORFs for accessory factors (9) (**Figure 1A**). The genome has a 5’ cap and 3’ polyA tail and serves initially as the template for translation of ORF1a and ORF1b, into the pp1a and pp1b polyprotein precursors that are processed by the viral proteases nsp3 and nsp5 to release 16 mature proteins (nsp1-16, key functions listed in **Figure 1A**) including all predicted components of the replication-transcription complex (RTC). The RTC produces negative strand (-) genomic and subgenomic (sg) RNAs from the genomic RNA. The generation of (-) sgRNAs is regulated by the transcriptional regulatory sequences (TRS). During the synthesis of the (-) RNA, the body TRSs (TRS-B) that precede each ORF (**Figure 1A**) act as potential dissociation signals for the RTC leading to a strand transfer to the TRS leader region of the 5’-UTR (TRS-L), to resume the synthesis of (-) sgRNA. The (-) RNAs are used as templates to synthesize full-length genomic RNAs and a nested set of sgRNAs. The frequency with which each TRS-B triggers dissociation during (-) RNA synthesis determines the relative abundance of each sgRNA and genomic RNA, such that the genomic RNA is relatively scarce while some sgRNAs, and consequently the proteins they code, are highly abundant (39).

**Figure 1.**
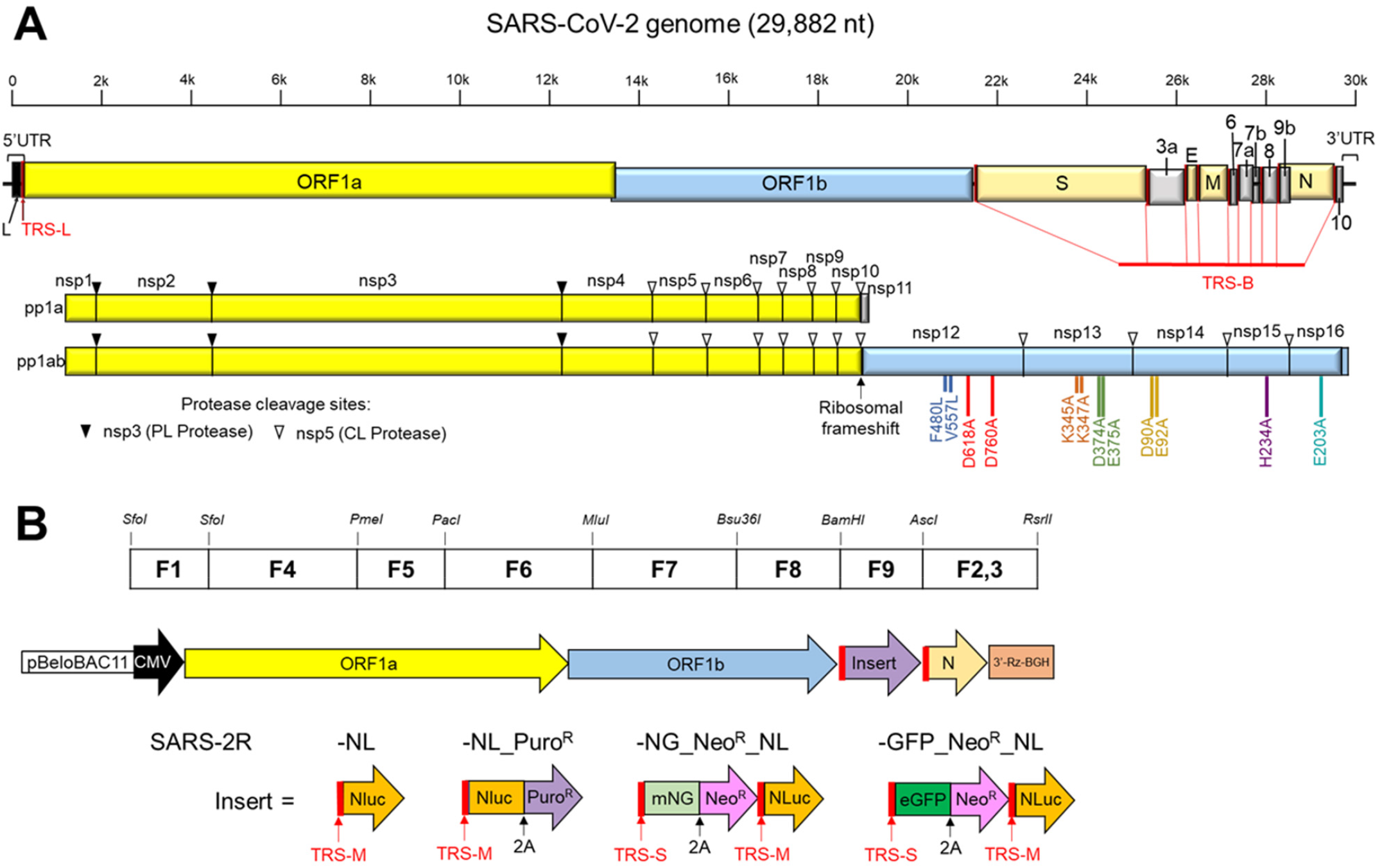
Genome organization of infectious SARS-CoV-2 and cloning strategy for the construction of the SARS-CoV-2 subgenomic replicon (SARS-2R). **A**. Genome structure of SARS-CoV-2 and open reading frames (ORF). L, is the leader sequence. Some non-structural proteins, nsp, and their functions are: nsp3, contains PLPpro, papain-like protease; nsp5, 3CLpro protease; nsp12, RNA-dependent RNA polymerase or RdRp; nsp13, helicase; nsp14, ExoN, exonuclease; nsp15, EndoU, endonuclease; nsp16, MTase, methyltransferase. PLPpro-, and 3CLpro-cleavage sites are depicted as black or white triangles, respectively. TRSs are transcription-regulatory sequences located immediately adjacent to ORFs; TRSs contain a conserved 6–7 nt core sequence (CS) surrounded by variable sequences. During negative-strand synthesis, nsp12 pauses when it crosses a TRS in the body (TRS-B) and switches the template to the TRS in the leader (TRS-L); UTR, untranslated regions. Select mutations in various replicons are shown: F480L/V557, blue-RDV-resistance mutations; D618A/D760A, red-nsp12 active site; K345A/K347A, brown-nsp13 nucleic acid binding; D374A/E375A, nsp13 active site; D90A/E92A light brown nsp14 active site; H234A, purple-nsp15 active site; E203A green-nsp16 active site. **B**. General strategy for the construction of SARS-CoV-2 replicons. Multi-step cloning strategy was used, whereby the fragments named SARS-CoV-2/F1 to SARS-CoV-2/F9 were sequentially cloned into pBeloBAC11 vector to generate pBAC-SARS-CoV-2-REP. The genetic structure of the replicon and the position of relevant restriction enzyme sites are shown at the ends of each fragment. F9 contains the reporter cassette. Early SARS-2R versions included T7 promoter (downstream from the CMV promoter) and were used for *in vitro* transcription of replicon RNA.

A variety of SARS-CoV-2 replicon systems have been published to date, featuring a range of advantages and disadvantages. Most involve the production of an RNA carrying one or more reporters and can be introduced into permissive cells by electroporation (24, 40-45). Some additionally have demonstrated the capability to be trans-encapsidated by co-expression with the deleted structural elements to produce single round infectious particles (41, 43, 45, 46).

Here, we report the generation of more than 20 SARS-CoV-2 replicons, including constructs containing orthogonal reporter proteins and selectable markers. The replicons are based on the “parental” Washington isolate (WA) as well as several <u>V</u>ariants <u>o</u>f <u>C</u>oncern (VOC) and <u>V</u>ariants <u>B</u>eing <u>M</u>onitored (VBM). We validate the replication dependence of reporter gene expression by use of replication-inactive mutants and well characterized inhibitors of viral enzymes, and thus the utility of these replicons for characterization of antiviral agents. We show that these subgenomic replicons can be delivered as plasmid DNA or transcribed RNA; we also show that the replicons can be trans-encapsidated by co-expression with SARS-CoV-2 structural proteins to produce virus-like particles (VLPs) capable of transducing angiotensin converting enzyme 2 (ACE2)-expressing cells (when bearing the SARS-CoV-2 S protein) or non-ACE2-expressing cells (when bearing the vesicular stomatitis virus glycoprotein). Importantly, we were able to generate stable cell lines harboring the SARS-CoV-2 replicon, a potentially valuable resource to facilitate identification and characterization of novel antivirals.

## RESULTS

### Construction of SARS-CoV-2 replicons (SARS-2Rs)

To facilitate the cloning of such large constructs, replicons were engineered in a bacterial artificial chromosomes (BAC) through stepwise cloning of DNA fragments (synthetic or prepared by RT-PCR from the SARS-CoV-2 Washington isolate, WA) into the BAC, exploiting unique restriction sites (**Figure 1B**). Many replicons have employed T7 promoters to permit *in vitro* transcription of replicon RNA, to be introduced into cells by electroporation (24, 40-45). To remove the necessity of generating RNA, an alternative approach employs a 5’ cytomegalovirus (CMV) promoter to drive initial transcription of replicon RNA (46-50). We built a replicon with a T7 promoter and a 3’ hepatitis delta virus ribozyme to ensure generation of the correct 3’ terminus. The base design incorporated the viral sequences essential for genome replication: the 5’ and 3’ UTRs, ORF1a and ORF1b, and N with its TRS-B. Reporters [Nano luciferase (NLuc) or NLuc with eGFP] and selectable markers (puromycin-N-acetyltransferase – Puro^R^ or neomycin phosphotransferase – Neo^R^) were inserted in place of M, E, S and most of the accessory ORFs, following the TRS-B sequences for M or S (TRS-M or -S) to permit transcription of the sgRNA. Placement of the reporter cassette in a sgRNA ensures that these genes will be expressed at a high level, but only following transcription by the SARS-CoV-2 RTC. This replicon produced a high luciferase signal that could be inhibited by RDV, indicating that the vast majority of reporter gene expression is generated by SARS-CoV-2 replication (**Figure S1**). However, to reduce labor and potentially increase reproducibility between constructs, we generated a second generation of replicon, replacing the T7 promoter with CMV promoter. After transfection of the BAC into target cells, transcription of full-length RNA is driven by the CMV promoter, allowing translation of ORF1a and ORF1b and formation of SARS-COV-2 RTC (**Figure 1**). The RTC then produces new gRNA and sgRNAs, *via* (-) RNA intermediates, leading to expression of the reporter genes and selectable markers. The BAC platform provides a stable platform for the generation of these large constructs; the introduction of unique restriction sites allows for modification of the reporter gene cassette or introduction of mutations into the replicon sequence through replacement of individual fragments. A partial list of replicons generated is shown in **Supplemental Table I**.

### Validation of replicon system in cell lines

To determine the optimal conditions for using nanoluciferase-producing SARS-CoV-2 replicon (SARS-2R_NL) we measured secreted NLuc over time in a range of cell lines (**Figure 2**). To differentiate between NLuc produced from the CMV promoter and replication-dependent NLuc expression, we used SARS-2R_NL with inactive nsp12 polymerase (nsp12_D618A/D760_). In all cases, nsp12_D618A/D760_ samples exhibited >99% decrease in NLuc activity, demonstrating that reporter gene expression is overwhelmingly replication dependent. We found maximum NLuc signal in 293T, BHK-21 and CHO-K1 cells, while the greatest signal-to-noise (WT ≥10,000-fold greater than nsp12_D618A/D760_) was achieved in BHK-21 and Caco2. As expected, mock controls (no SARS-2R_NL) produced no NLuc activity, while a robust signal for SARS-2R_NL WT over nsp12_D618A/D760_ was observed in all cell lines (**Figure 2**). In most cell lines, NLuc activity is measurable by 8 hours post transfection (hpt), with the signal typically plateauing 48 hpt. The differences in NL activity following SARS-2R_NL transfection among the various cell lines are likely dependent on a combination of transfection efficiency and differences in replication permissivity. These data demonstrate that protein expression from sgRNAs is replication dependent, and the reporter gene assays can be performed with high signal-to-noise in a range of convenient and/or physiologically relevant cell types, making this a suitable system for the study of SARS-COV-2 replication or screening for replication inhibitors.

**Figure 2.**
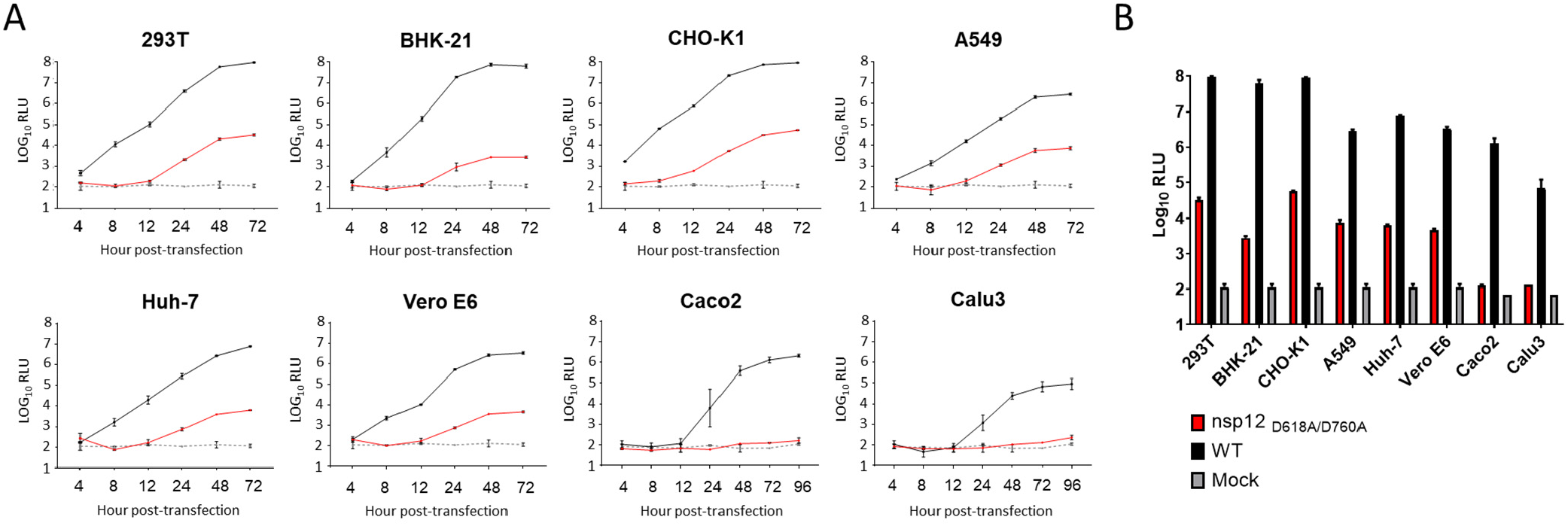
Time course of luciferase activity in various cell lines transfected with SARS-2Rs. **A**. NLuc activity kinetics for SARS-2R_NL replicons. Cells were seeded at 5×10^4^ cells/well in 24-well plates, 24 h before transfection. Cells were transfected with either mock (no plasmid) or 0.25 μg wild-type (WT) SARS-2R-NL or polymerase inactive SARS-2R-NL (nsp12_D618A/D760A_) plasmid and 0.025 μg N, ORF3b, ORF6 expression plasmids. NLuc was measured at the times indicated. **B**. Comparison of NLuc activity at 72 hpt. Average data from 2 independent experiments shown with standard deviation.

### Validation of replicase enzyme dependency of reporter gene expression

To determine whether SARS-2R replicons can be used to study the role of individual viral proteins and viral mutations in the replication mechanism of SARS-CoV-2 we constructed SARS-2R_NL_Puro^R^ replicons with mutations at various nsp active sites. Mutations at the predicted active sites of nsp12 (polymerase), nsp13 (helicase), and nsp15 (endonuclease), significantly reduce the activity of replicon activity in BHK-21 cells (**Figure 3A**). In particular, the nsp12_D618A/D760A_ and nsp13_D374A/E375A_ mutations entirely suppressed replication (>99.9% loss of activity). Similarly, the nsp15_H234A_ mutation resulted in >95% loss of activity. Mutations that were predicted to affect the nucleic acid binding function of SARS-CoV-2 helicase (51) (nsp13_K345A/K347A_) were also detrimental to replicon activity (>80% loss of activity). Interestingly, mutations at the exonuclease active site of nsp14 (nsp14_D90A/E92A_) and 2’-O-methyltransferase active site of nsp16 (nsp16_E203A_) only induced partial loss of replication (approximately 50% in each case), indicating that these enzymes/functions are less critical to replication, or that under these conditions the mutations may not cause a complete loss of activity.

**Figure 3.**
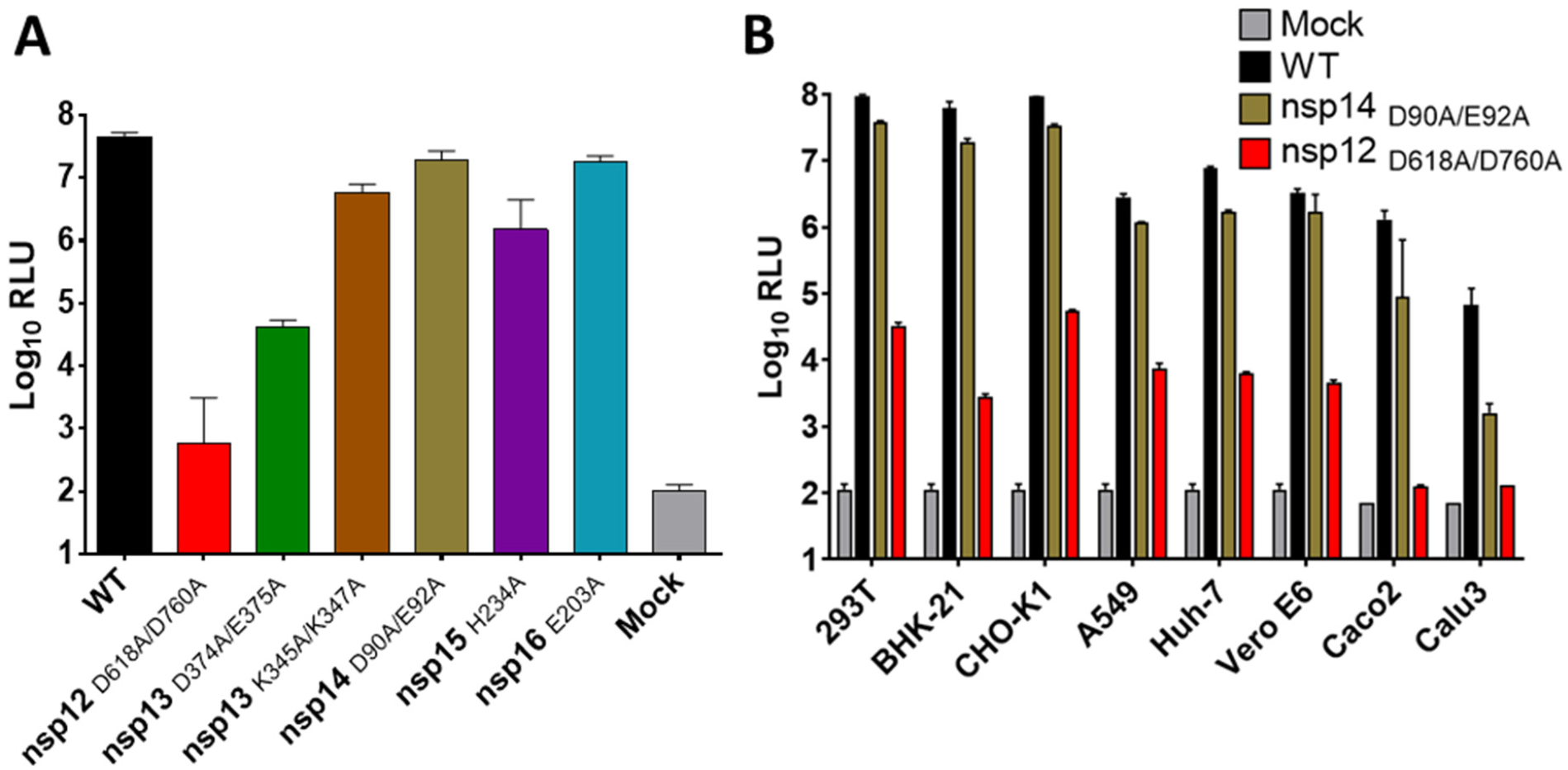
Fitness of SARS-2R-NLuc-Puro^R^ replicons bearing nsp mutations. Cells were transfected in 24-well plates with 0.25 μg of SARS-2R, together with 0.025 μg each of N-, ORF3b-, and ORF6-expression plasmids. Luciferase activity was measured at 48 hpt. **A**. Replicons carrying the indicated mutations were transfected into BHK-21 cells. **B**. Replicons that were either WT, ExoN(-) (nsp14_D90A/E92A_) or RdRp(-) nsp12_D618A/D760A_ were transfected into the indicated cell lines. Averaged data from 2 independent experiments are shown with standard deviations.

The nsp14 ExoN(-) mutant result was probed in more detail, as ExoN(-) mutations in nsp14 have been reported to impart variable replication defects in different betacoronaviruses, ranging from a moderate decrease in activity of the embecovirus Mouse Hepatitis Virus (MHV) (52) and sarbecovirus SARS-CoV (53) to essentially non-viable phenotype in merbecovirus MERS-CoV and the latest sarbecovirus SARS-CoV-2 (54). As more dramatic loss of replication fitness has been reported in nsp14 ExoN(-) mutant SARS-CoV-2 than we initially observed (54), we repeated our analysis of these mutants in a range of cells and consistently observed a moderate reduction in replicon activity (typically 50-75%); Calu3 cells exhibited a more profound reduction (∼98%) in activity with the ExoN(-) mutant replicon, however, replicon activity was lower generally in this cell line, potentially due to reduced transfection efficiency (**Figure 3B**). These data provide further validation of the replication-dependence of reporter gene expression, and indicate that this system would be particularly useful for studies of the polymerase and helicase, including screening for and characterization of small molecule inhibitors.

### Use of SARS-2Rs in studies of Variants of Concern (VOC) and Variants Being Monitored (VBM)

One of the major concerns during the COVID-19 pandemic has been the emergence of variants that exhibit properties such as faster spread, immune escape, and potentially increased pathogenicity. Although many studies have focused on the role of the S protein, there is increasing evidence that other mutations throughout the genome contribute to the phenotypic differences between variants. Additionally, mutations in the RTC enzymes have the potential to affect susceptibility to current and future therapeutics.

We generated SARS-2R_NG_Neo^R^_NL constructs derived from parental (WA), the alpha VBM B.1.1.7 (also known as “UK strain”), the beta VBM B.1.351 (also known as “South African or SA”), and the delta VOC B.1.617.2 (delta) viruses. Of note, the classification of SARS-CoV-2 variants as “of concern”, “of interest”, or “being monitored” has evolved during the course of the pandemic but the above variants have been thus classified by CDC as of 12/20/21. The VOC replicons differ from WA in ORF1a, ORF1b, N, and in non-coding regions (**Supplemental Table I**). To gain insight into whether the amino acid differences among variants in genes other than the S affect the replication efficiency of the B.1.1.7, B.1.351, and B.1.617.2 strains, we assessed replication in 293T cells (**Table I**). All replicons evinced clear replication above the polymerase-defective mutant WA replicon. However, the B.1.351- and B.1.1.7-derived replicons showed measurably reduced replication relative to the WA-derived replicon, suggesting that the mutations in these strains may affect fitness in the 293T cell culture model.

We next compared the antiviral susceptibilities of the VOC replicons in 293T cells. We found no statistically significant difference between the potency of RDV, NHC, EIDD-2801, and GC-376 among the four replicons (**Table I**). To verify that this lack of change represented the biology of the virus, we compared the *replicon* values for WA and B.1.1.7 to the values obtained with same compounds in fully infectious WA and B.1.1.7 viruses; indeed, we did not find significant differences in antiviral potency for any of the compounds against the two strains (**Supplemental Table II**). These data demonstrate the similarity between virus and replicon in terms of drug resistance, and that these constructs represent a valuable resource for comparative studies of drug susceptibility of the major variants.

**TABLE I.** Effect of Variants Being Monitored (VBM) and Variants of Concern (VOC) replicon mutations on replicon fitness and antiviral efficiency in 293T cells.

|  | **Fitness**<br>**(Fold change**<br>**relative to WT)** | **Mean EC<sub>50</sub> ± SD (μM)**<br>**(Fold change relative to WT)** |  |  |  |
| --- | --- | --- | --- | --- | --- |
| **Replicon** | **NLuc activity**<br>**(RLU)** | **RDV** | **NHC** | **EIDD-2801** | **GC-376** |
| **SARS-2R_NG_Neo<sup>R</sup>_NL**<br>**(WT or WA-derived)** | 2.7E+07 ± 8.3E+06<br>**(1.0)** | 0.004 ± 0.001<br>**(1.0)** | 0.75 ± 0.11<br>**(1.0)** | 4.92 ± 2.03<br>**(1.0)** | 0.36 ± 0.08<br>**(1.0)** |
| **SARS-2R_NG_Neo<sup>R</sup>_NL**<br>**B.1.1.7 (Alpha - VBM)** | 1.0E+07 ± 5.2E+06<br>**(0.4)** | 0.004 ± 0.001<br>**(1.0)** | 0.88 ± 0.16<br>**(1.2)** | 7.26 ± 1.49<br>**(1.5)** | 0.33 ± 0.03<br>**(0.9)** |
| **SARS-2R_NG_Neo<sup>R</sup>_NL**<br>**B.1.351 (Beta - VBM)** | 7.2E+06 ± 2.6E+06<br>**(0.3)** | 0.005 ± 0.001<br>**(1.3)** | 0.98 ± 0.10<br>**(1.3)** | 6.62 ± 0.80<br>**(0.7)** | 0.34 ± 0.03<br>**(0.9)** |
| **SARS-2R_NG_Neo<sup>R</sup>_NL**<br>**B.1.617.2 (Delta - VOC)** | 8.7E+07 ± 3.6E+07<br>**(3.2)\*** | 0.008 ± 0.001<br>**(2.0)\*** | 1.10 ± 0.05<br>**(1.5)** | 6.08 ± 0.62<br>**(1.2)** | 0.39 ± 0.07<br>**(1.1)** |

For all values N ≥ 2. Adjusted P Values in two-way ANOVA test comparing: Fitness of WT vs. Delta: 0.048; Alpha vs. Delta: 0.007; Beta vs. Delta: 0.007.

### Use of SARS-R2 in high-throughput assays

Initial experiments with SARS-CoV-2 replicons were conducted in 24-well plates. To provide evidence for the utility of SARS-2R as a tool for drug discovery efforts we explored conditions for its use in high throughput multi-well plate formats. We showed that reproducible measurements of RDV inhibition can be carried out in a 384-well format, at a range of cell seeding densities using 293T cells, without significant differences in the estimated EC_50_ and EC_90_s (**Table II** and **Figure S2**). Of note, these data demonstrate that the assay is reliable even at 1,000 cells/well, which is well within the range of an ultrahigh throughput format in 1536-well plates. Assay miniaturization is underway. These values are comparable to published data using infectious virus (20). The variation in RDV potency between cell types has been observed previously and likely reflects efficiency of conversion of RDV nucleotide to RDV-triphosphate (55, 56). Finally, to quantify the robustness of the assay, we calculated the Z’, comparing DMSO treatment to RDV; the average Z’ and 95 % CI was 0.70 ± 0.032. These data support the robustness and reproducibility of the replicon-based assay, and its suitability to high-throughput applications.

**Table II.** Inhibition of SARS-2R_NL_Puro^R^ by remdesivir based on dose response assays in a 384-well format using 293T cells.

| Cells/well | EC<sub>50</sub> (μM) | EC<sub>90</sub> (μM) |
| --- | --- | --- |
| 16,000 | 0.004± 0.001 | 0.028± 0.004 |
| 8,000 | 0.004± 0.001 | 0.018± 0.011 |
| 4,000 | 0.004± 0.001 | 0.011± 0.002 |
| 2,000 | 0.003± 0.001 | 0.013± 0.007 |
| 1,000 | 0.004± 0.002 | 0.012± 0.002 |

N = 2, standard deviation indicated.

We next used high throughput conditions with SARS-2R_NL_Puro^R^ to determine the potency of several antivirals with well-characterized activity against SARS-CoV-2 in 293T cells (**Table III**). Consistent with data reported for fully infectious SARS-CoV-2, the replicon was inhibited by RDV with an EC_50_ of 0.003 μM, EIDD-2801 with an EC_50_ of 4.67 μM, NHC with an EC_50_ of 0.76 μM, and by the nsp5 inhibitor GC-376 with an EC_50_ of 0.38 μM. We observed modest, but statistically significant (p<0.001) decrease in RDV susceptibility (3.3-fold), when comparing SARS-2R_NL_Puro^R^ WT to the replicon bearing the RDV resistance-associated mutations (nsp12_F480L/V557L_) (**Table III**) analogous to the modest resistance previously reported in MHV (nsp12_F476L/V553L_) and SARS-CoV (nsp12_F480L/V557L_) (up to 5.6-fold) (57). A decrease in RDV susceptibility was observed when these replicons were tested in BHK-21 cells, however, it was not found to be statistically significant under the testing conditions (not shown). The RDV-resistant replicon appears to retain susceptibility to NHC, EIDD-2801, and GC-376.

**TABLE III.** Inhibition of wild-type, RDV-resistant (nsp12_F480L/V557L_) and exonuclease inactive (nsp14_D90A/E92A_) SARS-2R_NL_Puro^R^ replicons by antivirals in 293T cells.

| Compound | EC<sub>50</sub> (μM) ± SD |  |  |
| --- | --- | --- | --- |
|  | WT | nsp12<sub>F480L/V557L</sub><br>(fold change from WT) | nsp14<sub>D90A/E92A</sub><br>(fold change from WT) |
| RDV | 0.003 ± 0.001 | 0.010 ± 0.005 **(3.3)\*** | 0.003 ± 0.001 **(1.0)** |
| NHC | 0.76 ± 0.12 | 0.62 ± 0.18 **(1.0)** | 0.73 ± 0.23 **(1.0)** |
| EIDD-2801 | 4.6 ± 2.4 | 4.5 ± 1.4 **(1.0)** | 5.0 ± 0.92 **(0.9)** |
| GC-376 | 0.38 ± 0.08 | 0.30 ± 0.04 **(1.3)** | 0.11 ± 0.02 **(3.5)\*** |

N=2, standard deviation indicated. Statistical significance indicated with \*. RDV, remdesivir; NHC, β-d-N4-hydroxycytidine; EIDD-2801, molnupiravir.

It is also of interest to understand whether inhibition by various antivirals that target nsp12 and are incorporated in the elongating RNA chain can be affected by the ExoN activity of nsp14. In MHV, the ExoN(-) mutant had increased sensitivity to RDV (57). Hence, we also tested the potency of such antivirals using the SARS-2R_NL_Puro^R^ nsp14_D90A/E92A_ replicon that lacks ExoN activity. We found that genetic suppression of this activity did not significantly affect the EC_50_ of NHC or EIDD-2801. Moreover, we did not observe any significant effect of the ExoN(-) mutations on RDV susceptibility (**Table III**); previously, Agostini et al. reported a 4.5-fold increase in sensitivity of the corresponding mutant MHV compared to the WT virus (57). Surprisingly, we did find a statistically significant increase in GC-376 susceptibility of SARS-2R_NL_Puro^R^ nsp14_D90A/E92A_ compared to WT (two-way ANOVA p-value 0.003). We are currently investigating whether (and how) the mutations that suppress the ExoN function of nsp14 affect the function of nsp5 and its interactions with GC-376.

In addition to resistance to an antiviral agent, the potential risk associated with drug-resistance mutations depends on their fitness. We used the replicon system to examine the effects of RDV-resistance associated mutations on fitness. Consistent with the reported modest decrease in production of infectious virus (57) our data show that the nsp12_F480L/V557L_ mutations reduce the replicon activity by ∼20 % (**Figure S3**). Interestingly, the nsp12_F480L/V557L_ mutations in the background of ExoN(-) nsp14 (nsp12_F480L_/_V557L_, nsp14_D90A/E92A_) seemed to significantly suppress replicon activity by 98%, far more than either pair of mutations alone. Collectively, these data illustrate the value of replicons for exploring the potential impacts of drug-resistance associated mutations on SARS-CoV-2 replication.

### Generation of SARS-CoV-2 replicon-expressing cell lines

While the use of replicon DNA to transfect BHK-21 cells streamlines the workflow relative to the use of transcribed RNA, the workflow can be simplified further through the use stable replicon harboring cell lines; similar tools have proven extremely beneficial in drug development for HCV (58, 59). Surprisingly, while a stable replicon harboring cell line was generated for SARS-CoV (60), none of the published SARS-CoV-2 replicon systems have led to the production of stable replicon harboring cell lines. We generated stable cells by transfecting BHK-21 cells with replicon DNA and then applying neomycin selection. GFP fluorescence and NLuc activity were monitored over time to confirm the presence of actively replicating replicons. After 4 weeks selection with neomycin clonal cell lines were selected and we identified one, BHK-SARS-2R_GFP_Neo^R^_NL, which exhibited GFP fluorescence and generated robust NLuc signal that persisted for multiple passages under selection and could be suppressed by treatment with RDV (>95%), indicating selection of a cell line with SARS-CoV-2 replication dependent reporter gene expression (**Figure 4**). Thus, this cell line can be used for multiplex assays that concomitantly follow NLuc and fluorescent signals, allowing for following highly sensitive and live replicon signal. There was no observable cytotoxicity due to the replicon and the cell morphology was typical of BHK-21 cells.

**Figure 4.**
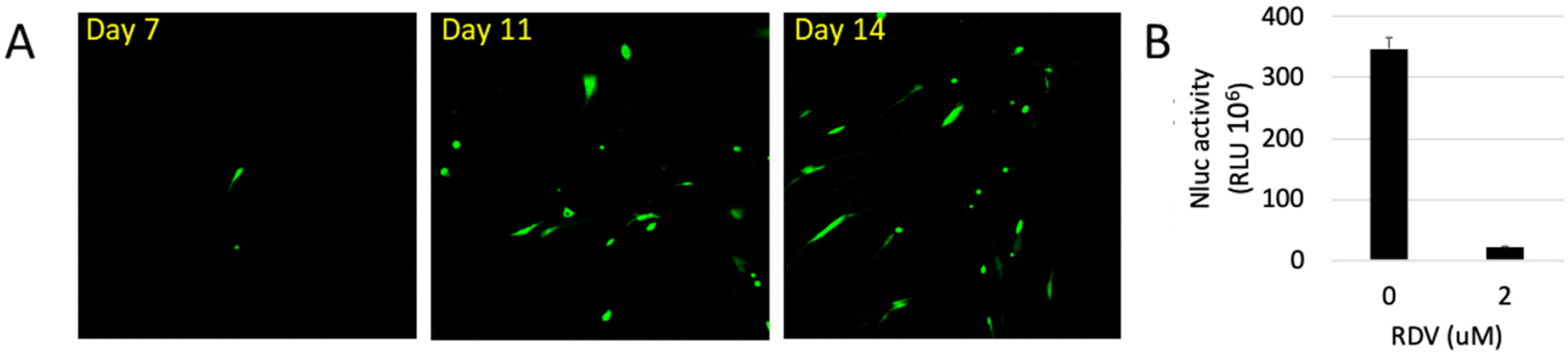
BHK-21 stable cell line of SARS-2 replicon. SARS-2R_GFP_NeoR_NL was transfected into BHK-21 cells in a 6-well plate. Cells were transferred to 10-cm dish 24 h post transfection and treated with 1 mg/ml G418 from 48 hpt. Colonies were picked and cultured under selection. **A**. Following establishment of a stable cell line GFP expression was monitored over time. **B**. NLuc activity was assayed 48 h following treatment with RDV.

The generation of a replicon stable cell line in which on a small percentage of cells exhibit evidence of active replicon gene expression is atypical and prompted further characterization. We performed qPCR and RT-qPCR to determine the levels of replicon RNA and plasmid DNA present in the cell line. After several weeks of passaging following transfection, we expected to find no replicon plasmid DNA, however, we found an estimated 1.2 (±0.9) replicon DNA copies per cell. We also found 35 (±26) replicon genomic RNA copies per cell. These data suggest that most/all of the cells carry an integrated copy of the plasmid, but at any given time only a small percentage of those cells contain active replicons.

To determine whether replicon activity was a stable phenotype, we separated the population of cells harboring active replicons (marked by GFP expression) from the overall population using fluorescence-activated cell sorting and acquired two populations of cells: GFP-positive (replicon active) and GFP-negative (replicon inactive). The phenotypes were confirmed by high content imaging and the cells cultured separately. The GFP-positive population rapidly declined from 100% to <5% of cells GFP positive after 48 h. Similarly, the proportion of GFP-positive cells in the GFP-negative sorted population climbed from 0% to almost 0.5% in 48 h (**Table IV**). After a week, the original phenotype of <5% GFP-positive (active replicon) was observed in both sorted populations, possibly because GFP-expression, and thus replicon activity, may not be a stable phenotype. The lack of replicon gene expression in most cells could be a consequence of inefficient transcription of the replicon RNA from the integrated DNA, or suppression of the replicon replication and gene expression by elements of the innate immune system. To better understand the replication characteristics of the BHK-SARS-2R_GFP_Neo^R^_NL cell line, we imaged live cells over 48 h in the presence of sodium butyrate (a histone deacetylase inhibitor) to relieve putative repression of transcription, ruxolitinib (a Janus kinase inhibitor) to block induction of IFN-induced gene expression, and RDV to block replicon activity (**Figure 5A, Figure S4 and Figure S5**). We found that sodium butyrate induced a 2-fold increase in GFP-positive cells relative to the untreated control, while ruxolitinib induced a strong 10-fold increase; treatment with both compounds was indistinguishable from ruxolitinib alone. Similar data were obtained using baricitinib, another JAK inhibitor (**Figure S6**). In all cases treatment with RDV prevented any increase in GFP-positive cells, confirming that the GFP-expression observed was dependent on the replicon RTC. Consistent with the hypothesis of unstable replicon replication, examination of the time-lapse images collected for the DMSO-treated BHK-SARS-2R_GFP_Neo^R^_NL cells revealed examples of GFP-positive cells appearing where there were previously none (**Figure 5B** – red arrowheads), a phenotype that was markedly more pronounced in ruxolitinib treated cells. There were also examples where cells that previously expressed GFP disappeared (**Figure 5B** – red circles), consistent with the reversion of the GFP-positive sorted cells to a predominantly GFP-negative population. Collectively, these data indicate that the replicon coding plasmid is integrated into the cell genome but is unable to establish persistent replication, consequently at any given time only a small percentage of cells contain active replicon replication. The time-lapse imaging results, in particular enhanced replicon activity in the presence of JAK inhibitors, suggests that the replicon stochastically activates but may be suppressed by IFN-mediated innate immune responses.

**Table IV.** Phenotypic stability of the BHK-SARS-2R_GFP_Neo^R^_NL replicon harboring cell line following sorting into homogeneous populations.

| Sorted population | % GFP positive cells |  |  |  |
| --- | --- | --- | --- | --- |
|  | 0 h | 24 h | 48 h | 168 h |
| GFP+ | 100 ± 0.0 | 12.7 ± 2.8 | 4.5 ± 1.3 | 0.95 ± 0.44 |
| GFP- | 0.0 ± 0.0 | 0.11 ± 0.03 | 0.43 ± 0.13 | 0.30 ± 0.05 |

N=2, standard deviation indicated.

**Figure 5.**
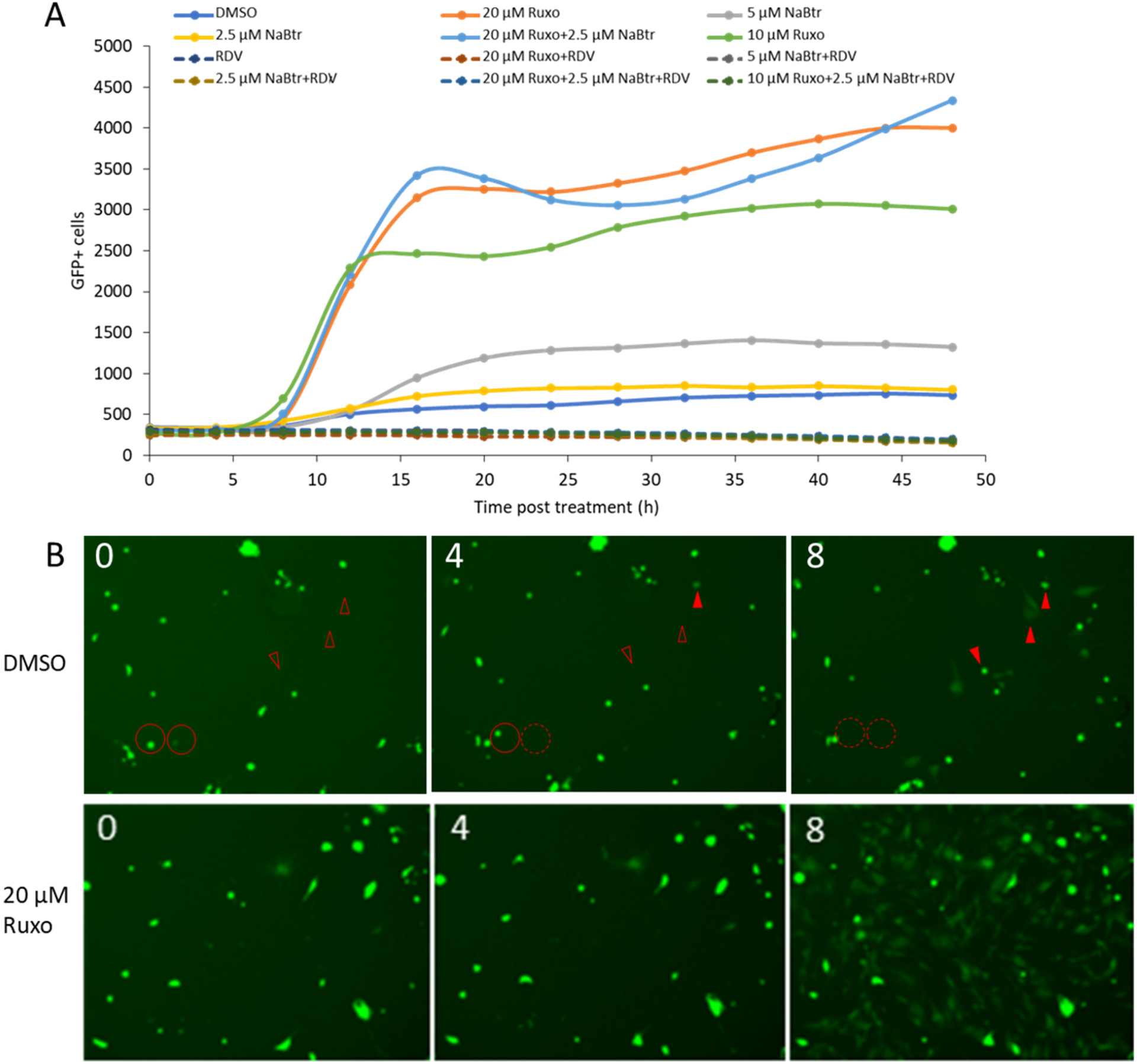
SARS-CoV-2 replicon activity following inhibition of IFN-signaling and derepression of transcription. BHK-SARS-2R_GFP_Neo^R^_NL cells were incubated with media containing the additives indicated. **A**. Replicon activity was assessed by counting GFP-positive cells. Images were collected at 4 h intervals using a high-content live-cell imaging system. **B**. Images of DMSO- and ruxolitinib-treated cells with time post-treatment indicated in hours. Cells with active replicon replication are GFP-positive. Solid red circles indicate active cells that disappear at later time points; dashed circles indicate the regions where the positive cells were. Solid red arrowheads indicate active cells that were not visible at earlier time points; empty red arrowheads indicate the same region prior to activation of the replicon. NaBtr, sodium butyrate; Ruxo, ruxolitinib; RDV, remdesivir. Representative data shown from one of two independent experiments.

### Validation of replicon-harboring cell lines for small molecule discovery and characterization

A major application for such a stable cell line is to facilitate compound screening and characterization of hit compounds. To validate the use of the BHK-SARS-2R_GFP_Neo^R^_NL cell line, we determined the EC_50_ for RDV using both the NLuc and GFP markers (**Table V and Figure S7**). The EC_50_ values determined using either reporter were generally comparable, both to one another and to values reported in the literature (2, 20, 27, 55). Of note, some differences in RDV potency have also been reported when tested in different cell lines, due to changes in its intracellular metabolism.

**Table V.** Determination of RDV efficacy using orthogonal reporter genes in the BHK-SARS-2R_GFP_Neo^R^_NL cell line.

|  | EC<sub>50</sub> (μM)<br>NLuc activity | EC<sub>50</sub> (μM)<br>GFP fluorescence |
| --- | --- | --- |
| RDV | 0.094 ± 0.001 | 0.082 ± 0.001 |
| NHC | 0.81 ± 0.18 | 0.88 ± 0.081 |
| EIDD-2801 | 6.0 ± 1.5 | 6.4 ± 2.8 |
| GC-376 | 0.29 ± 0.056 | 0.36 ± 0.042 |

N=2, standard deviation indicated.

### Generation of SARS-CoV-2 replicon-carrying VLPs by trans-encapsidation with SARS-CoV-2 structural proteins

The replicon system provides an invaluable resource for the study of processes related to replication of RNA and protein expression, however, the absence of essential structural proteins makes it impossible to study the processes of particle assembly and release, and virus entry. To expand the utility of the replicon system to address these functions, while retaining the advantages of a BSL2 compatible system, we supplied the M, E and S in trans by co-transfecting structural protein expression vectors with the replicon plasmid SARS-2R_GFP_Neo^R^_NL into 293T cells. After 48 h, media were collected and filtered, and used to transduce SARS-CoV-2 susceptible cell lines, either 293T-hAT or Huh7.5-hAT. After a further 48 h, reporter gene expression was measured as GFP-positive cells and NLuc activity in the media.

We were able to transduce both of the ACE2-expressing cell lines with VLPs bearing the SARS-CoV-2 spike protein (VLP_SARS2-S) or the SARS-CoV spike protein (VLP_SARS-S), but not with VLPs lacking any envelope protein (VLP_ Mock) (**Figure 6**). Comparing 293T cells (which do not endogenously express ACE2) to ACE2- and TMPRSS2-expressing 293T-hAT cells, we confirmed that both VLP_SARS2-S and VLP_SARS-S require ACE2 for successful transduction (**Figure 6A**). Similarly, comparing Huh7.5 cells to Huh7.5-hAT (Huh7.5 cells express ACE2; Huh7.5-hAT express additional ACE2 and TMPRSS2) revealed enhanced transduction with VLP_SARS2-S and VLP_SARS-S; by contrast VLP_ VSV-G particles were able to transduce the target cells; similarly, transduction of ACE2-expressing cells by VLP_SARS-2R_GFP_Neo^R^_NL_S was inhibited by a SARS-CoV-2 neutralizing monoclonal antibody. As a final control, we confirmed that reporter gene expression was dependent on replicon replication, following transduction, as all VLPs could be inhibited by treatment with RDV.

**Figure 6.**
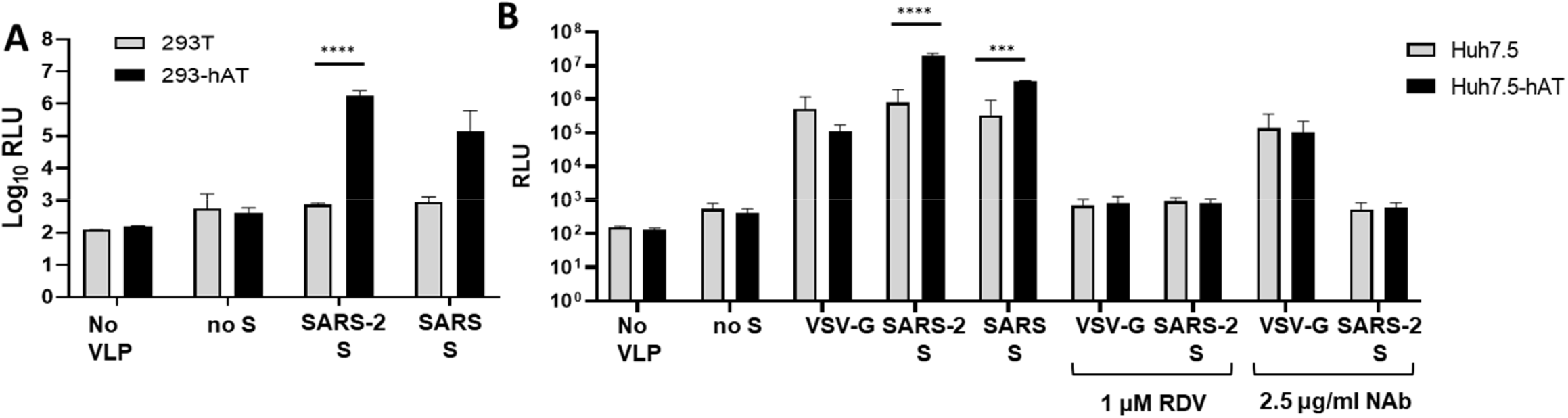
Trans-encapsidation of SARS-CoV-2 replicon. 293T cells were co-transfected with expression vectors for SARS-2R_GFP_Neo^R^_NL, E, M and one of SARS-2 S, SARS S VSV-G or empty vector. After 48 h, VLP media were collected, filtered and used to transduce the cell lines indicated. NLuc activity was measured 48 h post transduction. **A**. Transduction of 293T and 293T-hAT with VLPs bearing SARS S or SARS-2 S. **B**. Transduction of Huh7.5 and Huh7.5-hAT with VLPs bearing SARS S, SARS-2 S or VSV-G. Prior to transduction, some cells were treated with RDV to inhibit replication, or an S-targeting neutralizing antibody (NAb) to prevent VLP entry. P values for pairwise comparisons determined by 2-way ANOVA: ****, p ≤ 0.0001; *** p ≤ 0.001. Means shown with standard deviation. N=3, except for “No VLP” and “VSV-G” where N=2.

## DISCUSSION

Despite significant success in the fight against COVID-19, both in the development of effective vaccines and some small molecule therapeutics, SARS-CoV-2 has proven remarkably resistant to elimination efforts. The persistence of the global pandemic and the continuing evolution of the virus make it ever more important to expand research into both coronavirus biology and SARS-CoV-2 therapeutics specifically. A primary limitation in SARS-CoV-2 research is the high risk associated with working with a pathogenic respiratory pathogen, in terms of both the availability of suitable facilities and of suitably trained researchers. Here we present a range of approaches to facilitate research into all facets of SARS-CoV-2 replication, including cell entry, nucleic acid transcription and replication, and virus assembly and release; collectively these systems are tools to identify and characterize inhibitors at any point in SARS-CoV-2 replication. These systems are based on subgenomic SARS-CoV-2 replicons carrying reporter genes, reducing biological hazards and providing means to easily measure viral replication.

The most popular strategy for generating replicons from positive-strand RNA virus is to transcribe RNA *in vitro* that can be electroporated into target cells to initiate replication. To facilitate use of the assay, particularly in the context of large-scale screening, we eliminated the need for *in vitro* transcription by cloning cDNA for the SARS-CoV-2 replicons into a BAC under control of a CMV promoter. The replicon comprised all replication-essential proteins and sequences but omitted the structural proteins S, E, and M; additionally, reporter genes were included as a separate ORF using the TRS-B of M to drive production of the sgRNA expressing the reporter cassette. Placing the reporter cassette in a sgRNA ensured that formation of a functional RTC was essential for reporter gene expression. We validated the replication dependence using both genetic ablation of essential viral enzymes and the use of well characterized inhibitors of SARS-CoV-2 replication. Combining these controls with a screen of suitable cell lines, we were able to demonstrate replication-dependent reporter expression with signal-to-noise ratios between 1:100 and 1:>10,000; this wide dynamic range combined with an excellent z’ score indicate that these replicons will be well suited to small molecule screening and should be able to resolve a wide range of magnitude of phenotypes associated with modulation of viral or cellular factors. Additionally, we built replicons based not only on the early WA1 isolate, but also more recent variants, including alpha VBM B.1.1.7, beta VBM B.1.351, delta VOC B.1.617.2, and epsilon VBM CAL.20C (characterization of epsilon and cloning of omicron are in progress). Liu et al. highlighted that from May 2 to July 31 of 2021, the prevalence of the delta variant in the USA increased (from ∼1 % to 94 %), while that of the alpha variant decreased (from 70% to 2.4%) (61). They also showed that the delta P681R mutation on S enhanced the cleavage of the full-length spike to S1 and S2, leading to increased infection *via* cell surface entry. The measurably increased replication of the delta replicon suggests that the increased transmissibility of the delta VOC may also be due in part to mutations at regions outside the Spike protein. This hypothesis is currently under investigation.

In addition to the dynamic range of the assay, a major concern when screening for small molecules inhibitors of viral replication is the potential of scoring inhibitors of the reporter gene as antiviral hits. To mitigate that risk, we have generated replicons with orthogonal reporter genes (NLuc and GFP), an approach successfully employed by others (40, 46, 50), and shown that measurements of antiviral potency are similar, whichever reporter is used.

In a final step to enhance the utility of these replicons for small molecule screening, we generated a BHK-21-based stable cell line harboring the dual-reporter replicon and showed that it can be used to determine the potency of antiviral compounds. Stable cell lines are of particular use for high throughput screening, to this end we demonstrated the utility of this stable cell line for high-throughput screening by determining the potency of RDV *vs*. SARS-CoV-2 in a 384-well plate format. The combination of orthogonal reporters, high sensitivity for use in 384-well plates, and the convenience of the stable cell line format, combined with the biological safety of a replicon (most robotic screening facilities are not located in BSL3 laboratories), make this system highly suitable for small molecule discovery and characterization efforts.

While not diminishing the utility of this stable cell line, it is worth noting that it differs from the majority of replicon-harboring cell lines, in that the host cells contain replicon-coding DNA integrated into their genome. While most replicon-harboring cell lines are based on self-replicating RNA expressing a selectable reporter (35, 60), this approach has proven challenging for SARS-CoV-2 as we and others have explicitly tried and failed to generate such cell lines. Possible explanations include potential suppression by IFN-mediated immune responses and/or killing of replicon-harboring cells by expression of viral proteins such as nsp1 (50, 62, 63). Indeed, a recent pre-print reports the generation of a mutant replicon-harboring cell line, with a mutation in nsp1 (64). These factors may contribute to the replicon quiescence we observed in the majority of cells and the lack of sustained reporter gene expression following initiation of replication. Significantly, when the cell line was sorted for homogeneous populations of reporter gene expressing or non-expressing cells, we found that both rapidly reverted to the pre-sorted proportion of cells harboring active replication. This suggests that the cells in the cell line may be continuously initiating transient replication but failing to sustain replicon replication. It may also be relevant that the cell line could be established with G418 selection but not puromycin. If cells are cycling between active and inactive states, it may be that the longer time to induce cell death with G418 is necessary to give the cells the opportunity to produce neomycin phosphotransferase, the more rapid toxicity of puromycin may kill cells when the replicon is inactive.

If the replicon-harboring cells are only transiently exhibiting RNA replication, this system may be unsuited for evolution of resistance studies, as the replicons will not have the opportunity to acquire mutations and persist over time under selection; nevertheless, our approach provides a powerful, convenient and reproducible system for library screening and studies of putative antivirals. Moreover, reproducibility will be enhanced by continuous re-initiation of replicon expression from the DNA, such that the replicon will not deviate from the cloned sequence over time; in autonomously replicating RNA cell lines, replicons are often seen to undergo culture adaptation (for example (64, 65)). The stability of these DNA-based replicon-harboring cell lines could be of particular value when it is necessary to be able to compare potencies of compounds over large numbers of experiments.

Finally, to expand the utility of the replicons in the study of SARS-CoV-2 replication, we complemented replicon expression with expression of structural proteins, as has been employed in other RNA virus replicon systems (66) including SARS-CoV-2 (41, 43, 45, 46), and were able to demonstrate transmission of SARS-CoV-2 particles bearing S (of either SARS-CoV or SARS-CoV-2) or VSV-G. Reporter gene expression was shown to be dependent on receptor binding and replication, making this a viable approach to study effects on particle assembly and release, and binding and fusion to target cells. Introduction of these replicons with cells expressing structural proteins *in trans* may permit viral spread and thus passaging, as seen in other SARS-CoV-2 replicon systems (41, 45).

In conclusion, the replicon-based systems described here provide tools to examine all facets of the SARS-CoV-2 replication cycle, from cell entry, through protein expression and genome replication, to particle assembly, and can be applied to the diverse range of existing and emerging virus variants. In addition to the numerous applications to basic research, the production of genetically stable, cDNA-based, replicon-harboring cell lines has great potential value for the identification and evaluation of antivirals. The use of integrated cDNA-based replicons may also represent a viable strategy for the production of stable cell lines based on RNA virus replicons that do not readily establish persistent RNA-based replicon-harboring cell lines.

## MATERIALS AND METHODS

### Cells and viruses

VeroE6 (ATCC) are derived from African green monkey kidney cells and lack the genes encoding type I interferons (67, 68). Huh-7.5 (provided by Charles Rice) are derived from human hepatoma cells. Huh7.5-hAT cells (provided by Gregory Melikian and Mariana Marin), have been modified to stably express human ACE2 and human TMPRSS2, to enhance infection by SARS-CoV-2. A549 (ATCC) are derived from human adenocarcinoma cells. Human embryonic kidney (HEK) 293T cells stably express the SV40 large T antigen, are resistant to neomycin, and were selected for their high transfectability (ATCC). 293T-hAT cells (provided by Bruce Torbett) have been modified to stably express human ACE2 and human TMPRSS2, to permit infection by SARS-CoV-2. Baby hamster kidney (BHK)-21 are a popular substrate for viral growth and may have defects in IFN production and response to IFN (69, 70). These cells were all cultured in humidified conditions at 37 °C in 5% CO_2_ in Dulbecco’s modified Eagle’s medium (DMEM) supplemented with 10% fetal bovine serum (FBS), 2 mM L-glutamine, non-essential amino acids (NEAA), and 100 U/ml penicillin and 100 μg/ml streptomycin.

Chinese hamster ovary (CHO)-K1 cells (ATCC) are a clonal derivative of the parental CHO cell line and were cultured in humidified conditions at 37 °C in 5% CO_2_ in F12 medium supplemented with 10% FBS, 2 mM L-glutamine, and 100 U/ml penicillin and 100 μg/ml streptomycin.

Caco2 cells (ATCC) are a heterogeneous population derived from human colorectal adenocarcinoma and can reflect a wide variety of cell types and properties found in primary tissue. Calu3 cells (ATCC) are a heterogeneous population derived from human lung adenocarcinoma tissue. Both Caco2 and Calu3 cells were cultured in humidified conditions at 37 °C in 5% CO_2_ in Eagle’s minimum essential medium (EMEM) supplemented with 10% FBS, 2 mM L-glutamine, NEAA, and 100 U/ml penicillin and 100 μg/ml streptomycin.

SARS-CoV-2 strains 2019-nCoV/USA_WA1/2020 and SARS2 hCoV-19/England/204820464/2020 (B.1.1.7) were from BEI Resources and represent the parental and alpha lineages of SARS-CoV-2.

### Plasmids, antivirals and chemicals

Plasmids pLVX-EF1a-IRES-Puro SARS-CoV-2 E_NR-52967 and pLVX-EF1a-IRES-Puro SARS-CoV-2 M_NR-52968 (BEI) were gifts from Nevan Krogan (University of California, San Francisco) (71). For our purposes, the E and M coding regions of each plasmid were amplified by PCR, without the STREP tags, and re-cloned into the vector using the NEBuilder kit (New England Biolabs) following the manufacturer’s instructions. SARS-S in pCAGGS was a gift from Paul Bates (University of Pennsylvania) (72). SARS-CoV-2 S was synthesized (Genscript) and cloned into pcDNA3.1 between two PmeI sites using NEBuilder. The plasmid sequence was confirmed by Sanger sequencing. The replicons were cloned into pBeloBAC11 (NEB) (73). The compounds RDV (#30354), EIDD-1931 (#9002958), EIDD-2801/molnupiravir (#29586), GC-376 (#31469), ruxolitinib (#11609), and sodium butyrate (#13121) were purchased from Cayman Chemicals. A chimeric neutralizing antibody specific against SARS-CoV-2 S protein receptor binding domain (NR-55410) was provided by ACROBiosystems through BEI resources.

### Construction of the SARS-CoV-2 subgenomic replicon (SARS-2R)

DNA fragments of SARS-CoV-2 replicon were either synthesized from Integrated DNA Technologies Inc or amplified by reverse transcription and PCR (RT-PCR) with viral RNA template that was extracted from the supernatants of SARS-CoV-2-infected Vero cells (SARS-CoV strain 2019-nCoV/USA_WA1/2020 (WA1). RT was performed by using the SuperScript® III First-Strand Synthesis System (ThermoFisher Scientific, Catalogue number 18080051) with gene specific primers (SEM296 5’-ctttaattaactgcttcaacc-3’, SEM297 5’-tagtgcaacaggactaagctc-3’, SEM298 5’-tgaagtctgtgaattgcaaag-3’, SEM299 5’-acccagtgattaccttactac-3’, SEM300 5’-gttacagttccaattgtgaag-3’) and oligo dT primer. Multi-step cloning strategy was used, whereby the fragments named SARS-CoV-2/1 to SARS-CoV-2/9 were sequentially cloned into pBeloBAC11 vector to generate pBAC-SARS-CoV-2-REP. Insertions were performed using the NEBuilder kit following the manufacturer’s instructions. The genetic structure of the replicon and the position of relevant restriction enzyme sites are shown at the ends of each fragment. F9 represents the region of the reporter genes.

The sequences of the SARS-2R replicons were confirmed by both Sanger and deep sequencing (initial WT constructs), and subsequent mutations were confirmed by Sanger sequencing the relevant regions.

### T7 transcription and RNA electroporation

The DNA template for T7 transcription was prepared by linearizing the T7 replicon plasmid (#453, Supplemental Table I) with the restriction enzyme SacII (NEB), followed by protease K treatment, phenol/chloroform extraction and ethanol precipitation. Replicon RNA was *in vitro* transcribed with the mMESSAGE mMACHINE T7 Transcription Kit (ThermoFisher Scientific) according to the manufacturer’s instructions. An 80 μl reaction was set up by adding 8 μg linearized DNA template and 12 μl GTP (cap analog-to-GTP ratio of 1:1). The reaction mix was incubated at 32 °C for 12 h. Following transcription, template DNA was removed with Turbo DNase, and RNA was recovered with phenol/chloroform extraction and isopropanol precipitation as suggested by the manufacturer. A second transcript coding for SARS-CoV-2 nucleocapsid protein N was *in vitro* transcribed from a PCR product (forward 5’-TACTGTAATACGACTCACTATAGGatgtctgataatggaccccaaaatc-3’ [T7 promoter in uppercase] and reverse 5’-TTTTTTTTTTTTTTTTTTTTTTTTTTTTTTTTTTTTaggcctgagttgagtcagcac-3’ primers) using the mMESSAGE mMACHINE T7 Transcription Kit. A 20 μl reaction was set up by adding 0.2 μg of N PCR product and 1 μl GTP (cap analog-to-GTP ratio of 2:1). The reaction was incubated at 37 °C for 4 h. N mRNA was purified with RNeasy mini kit (Qiagen).

BHK-21 and 293T cells were trypsinized and washed twice with DPBS (Sigma-Aldrich). BHK-21 cells were adjusted to 1 × 10^7^ cells/ml and 293T cells were adjusted to 5 × 10^6^ cells/ml. For each electroporation, 250 μl cells were pelleted and resuspended in 250 μl of Ingenio electroporation solution (Mirus). For each electroporation, 2.5 μg of each replicon RNA and N mRNA were mixed with cells and transferred to a 0.4 cm electroporation cuvette (pre-chilled on ice). Cells were pulsed once at 280 V for BHK-21 cells and 250 V for 293T cells with Ingenio EZporator MIR51000 electroporation system in low voltage (LV) mode (150 Ω internal resistance and 1050 μF capacitance). After 5 minutes recovery at room temperature, the electroporated cells were transferred to a 10 cm dish or seeded into a 96-well plate. Nano luciferase assays were performed 48 h post electroporation.

### Comparison of replicon fitness by transient transfection

Cells were seeded in 24-well plates and incubated until approximately 80 percent confluent. Then cells were transfected with either mock (no replicon) or 0.25 μg replicon plasmid and 0.025 μg N, ORF3b, ORF6 expression plasmids with jetPRIME transfection reagent (Polyplus). In all cases, the amount of SARS-2R nucleic acid transfected was quantified by both agarose gel electrophoresis with ethidium bromide staining and quantification of the absorbance at 260 nm and 280 nm using a NanoDrop spectrophotometer (ThermoFisher Scientific). Nano luciferase activity was assayed using the Nano-Glo Luciferase Assay System (Promega) following manufacturer’s instructions. The luciferase signal was measured using a GloMax Navigator Microplate Luminometer (Promega).

### Determination of Replicon EC_50_ and EC_90_ values

BHK21 cells seeded in 6-well plate were transfected with SARS-2R using jetPRIME transfection reagent (Polyplus transfection). At 16 h post transfection, cells were trypsinized then seeded into 96- or 384-well plate and treated with serial diluted compounds. Stable cells were seeded directly and treated with serial diluted compounds. Nano luciferase assays were performed 48 hours post treatment. Dose response curves were calculated with Prism software.

### Generation of cells that carry SARS-2R replicon

BHK-21 cells were seeded into a 6-well plate, then transfected with replicon SARS-2R_NG_Neo^R^_NL using jetPRIME transfection reagent following the manufacturer’s instructions. The following day, cells were trypsinized and transferred into a 10 cm dish. After a further 24 h incubation, G418 was added at 1 mg/ml and cells were maintained until distinct colonies formed. Individual colonies were picked and transferred to a 24-well plate. These clones were expanded until near confluency, then NLuc activity was assayed to confirm the presence of active replicon replication. Clones that exhibited a robust NLuc signal were trypsinized then seeded into two separate wells, one treated with 1 μM RDV to inhibit replicon replication. Clones that exhibited a robust NLuc signal that could be inhibited with RDV were expanded for future studies.

### Flow cytometry and subsequent imaging

BHK-SARS-2R_GFP_NeoR_NL cells were trypsinized and washed twice with DPBS (Sigma-Aldrich) and once with sort buffer (PBS supplemented with 0.3 % BSA, 2 mM EDTA, 25 mM HEPES pH 7.0). Cells were adjusted to 1-2 × 10^7^ cells/ml in sort buffer. GFP+ and GFP-cell sorting was carried out using a BD FACS Aria II SORP Cell Sorter (Pediatric/Winship Flow Cytometry Core, Emory University). After sorting, GFP+ and GFP-cells were seeded into 96-wells at 2500-5000 cells/well. Cells were stained with Hoechst 33342 and counted for GFP+ and total cells with Cytation 5 high-content live-cell imaging system (Biotek) using a 4x objective.

### Live cell imaging of replicon harboring cell lines following modulation of cellular transcription

Cells were seeded into 96-well plates at 10,000 cells per well. After 24 h, culture media were removed and replaced with media supplemented with ruxolitinib suppress IFN signaling, sodium butyrate to enhance transcription, and/or RDV to suppress SARS-CoV-2 replication. GFP-positive cells were imaged on a Cytation 5 high-content live-cell imaging system (Biotek) using a 4x objective at 37 °C in 5% CO_2_ over 48 to 72 h. Images were captured as a 5×4 matrix and GFP positive cells were enumerated using Gen5 software (Biotek).

### qPCR to determine levels of SARS-CoV-2 nucleic acid

RNA and DNA was harvested from the BHK-SARS-2R_GFP_Neo^R^_NL cell line using the RNeasy mini kit (Qiagen) and the QIAamp genomic DNA mini kit (Qiagen), respectively. RNA was treated with DNase I (Qiagen) before reverse transcription. Reverse transcription was performed by using the SuperScript® III First-Strand Synthesis System (ThermoFisher Scientific) with a gene specific primer 5’-tgaagtctgtgaattgcaaag-3’. qPCR was performed with QuantStudio 3 and ABsolute qPCR SYBR Green Mix (Thermo Fisher Scientific) using primers SEM397 5’-cttatgattgaacggttcgtgtc-3’ and SEM398 5’-cagaatacatgtctaacatgtgtcc-3’. SARS-CoV-2 nucleic acid copy numbers were derived from a calibration curve comprising six 10-fold dilutions of plasmid DNA for SARS-2R_GFP_Neo^R^_NL. Final values were averaged from two independent experiments and shown with 95% confidence intervals.

### Production of SARS-CoV-2 stocks

SARS-CoV-2 stocks were produced by infection of VeroE6 with SARS-CoV strain 2019-nCoV/USA_WA1/2020 or SARS2 hCoV-19/England/204820464/2020 (B.1.1.7). Media were harvested after 48 hours or when cytopathic effect was observed. To titer viral stocks, VeroE6 cells were seeded in a 96-well plate at 20,000 cells per well. After 24 h, infections were performed with serially diluted viral stocks. Cells were fixed 6 hours post infection (i.e., long enough for expression of viral proteins but short enough that no spread will have occurred) and stained for SARS-CoV-2 N protein to identify infected cells. Infected and total cells were determined by high content microscopy, virus titer was calculated as the number of infected cells per volume of virus stock added to cells.

### Determination of SARS-CoV-2 EC_50_ values

Calu3 cells were seeded in a 96-well plate at 30,000 cells per well and infected when at least 80% confluent. Prior to infection, medium was replaced with fresh medium and supplemented with serially diluted compounds. Duplicate wells for each condition were infected at an MOI of 0.01, fixed 48 h post infection and stained for N protein expression. The number of infected cells at each compound concentration was determined by high content microscopy and dose response curves were plotted and EC_50_s calculated with Prism software.

### Immunofluorescent staining

SARS-CoV-2 infected cells were fixed in 96-well plates with 4% paraformaldehyde in phosphate buffered saline (PBS) for 30 min at room temperature, then incubated in PBS supplemented with 0.1% Triton X-100 for 5 min at room temperature, followed by incubation in blocking buffer (5% FBS in PBS) for 1 h at room temperature. Rabbit monoclonal anti-N antibody (Sino Biologicals #40143-R001) was diluted 1:5,000 in 0.1% Tween-20 in PBS (PBS-T), and incubated on samples overnight at 4 °C. Samples were washed twice in PBS-T, then incubated in secondary goat anti-rabbit Alexa Fluor 647 (Invitrogen #A-21244) diluted 1:2,000 and Hoecsht-33342 (ThermoFisher, MA, USA) at 0.5 μg/ml in PBS-T, for 1 h at room temperature. Samples were washed four times in PBS-T. Imaging was performed using a Cytation 5 multimode imaging system with Gen 5 software (Biotek).

### Preparation of VLPs

293T cells were seeded into a 6-well plate at 1.2 × 10^6^ cells/well then incubated at 37 °C for 24 h. Each well was transfected with 1 μg replicon SARS-2R_GFP_NeoR_NL, and 0.25 μg each of N, E, and M expression vectors, and 0.25 μg of expression vectors for one of SARS-CoV S, SARS-CoV-2 S, VSV-G or an empty vector, with 4 μl jetPRIME transfection reagent (Polyplus). VLPs were collected 48 h post transfection and passed through a 0.45 μM filter. A 96-well plate was treated with 20 μg/ml poly-D-Lysine (Sigma-Aldrich) for 1h at room temperature and washed twice with 1 x DPBS (Sigma-Aldrich). Target cells were seeded at 4 × 10^4^ cells/well (293T, 293T-hAT) or 2 × 10^4^ cells/well (Huh-7.5, Huh7.5-hAT) and incubated at 37 °C for 24 h. Target cells were infected with 50 μl VLPs per well. For antibody treatment, VLPs were mixed with 2.5 μg/ml neutralizing antibody (BEI NR-55410, provided by ACROBiosystem, cat # SPD-M128) and incubated at 37 °C for 1 h before adding to the target cells. Cells were washed 3 times with PBS at 24 h post infection, then incubated in DMEM for a further 24 h. Nano luciferase assays were performed 48 h post infection.

### Calculation of Z’

Cells were seeded in a 6-well plate then transfected with SARS-2R and treated with DMSO or RDV. After 48 h, the NLuc activity in the media was measured for 3 replicate samples per condition. The mean (μ) and standard deviation (σ) were calculated for each treatment condition. Z’ was calculated as 1-[3σ DMSO + 3σ RDV]/[μ DMSO - μ RDV]; an excellent assay should have Z’ > 0.5, indicating a wide range between positive and negative controls, combined with small standard deviations. A theoretically perfect assay would have Z’ = 1.

### Statistics

Data were tabulated and graphs plotted using Excel (Microsoft) or Prism (Graphpad). Statistical tests were performed using Prism.

## ACKNOWLEDGEMENTS

This research was supported in part by the Nahmias-Schinazi Chair fund at Emory University (SGS). The following reagents were obtained through BEI Resources, NIAID, NIH: SARS-CoV-2 strains 2019-nCoV/USA_WA1/2020 and SARS2 hCoV-19/England/204820464/2020 (B.1.1.7); monoclonal Anti-SARS-Related Coronavirus 2 Spike Glycoprotein Receptor Binding Domain (RBD), Chimeric Potent Neutralizing Antibody (produced in vitro), NR-55410. We wish to thank Nevan Krogan for plasmids pLVX-EF1a-IRES-Puro SARS-CoV-2 E_NR-52967 and pLVX-EF1a-IRES-Puro SARS-CoV-2 M_NR-52968 and Luis Enjuanes for the pBeloBAC11 plasmid. Also Raymond F. Schinazi and Keivan Zandi, for the stock of the WA1 strain and to Ann Chahroudi and Nils Schoof for the stock of B.1.1.7 strain. We also thank Charles M. Rice (Rockefeller University), for the Huh7.5 cells, Gregory Melikian and Mariana Marin (Emory University) for the Huh7.5-hAT cells, Bruce E. Torbett (University of Washington) for the 293T-hAT cell, and Eleftherios Michailidis and Raymond F. Schinazi for useful discussions. We acknowledge the Pediatric/Winship Flow Cytometry Core, Emory University and its personnel for help with use of the BD FACS Aria II SORP Cell Sorter. Finally, we are grateful to Jindi Lan (deceased) for his steadfast support of Shuiyun Lan and family, enabling this research to continue during the COVID-19 pandemic and shutdowns.

## Supplemental Tables and Figures

**SUPPLEMENTAL TABLE I.**
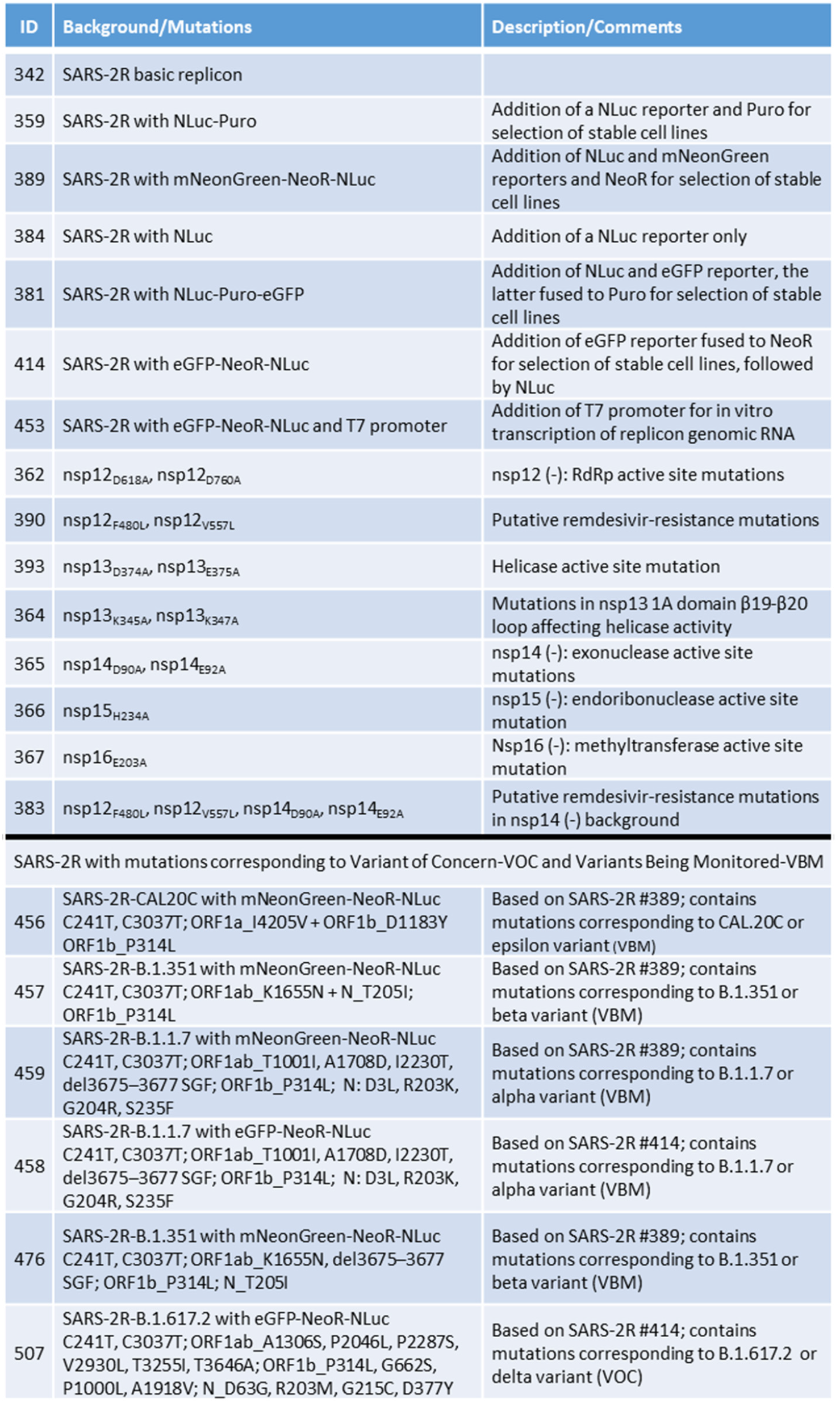
List of constructed SARS-2Rs.

**SUPPLEMENTAL TABLE II.**
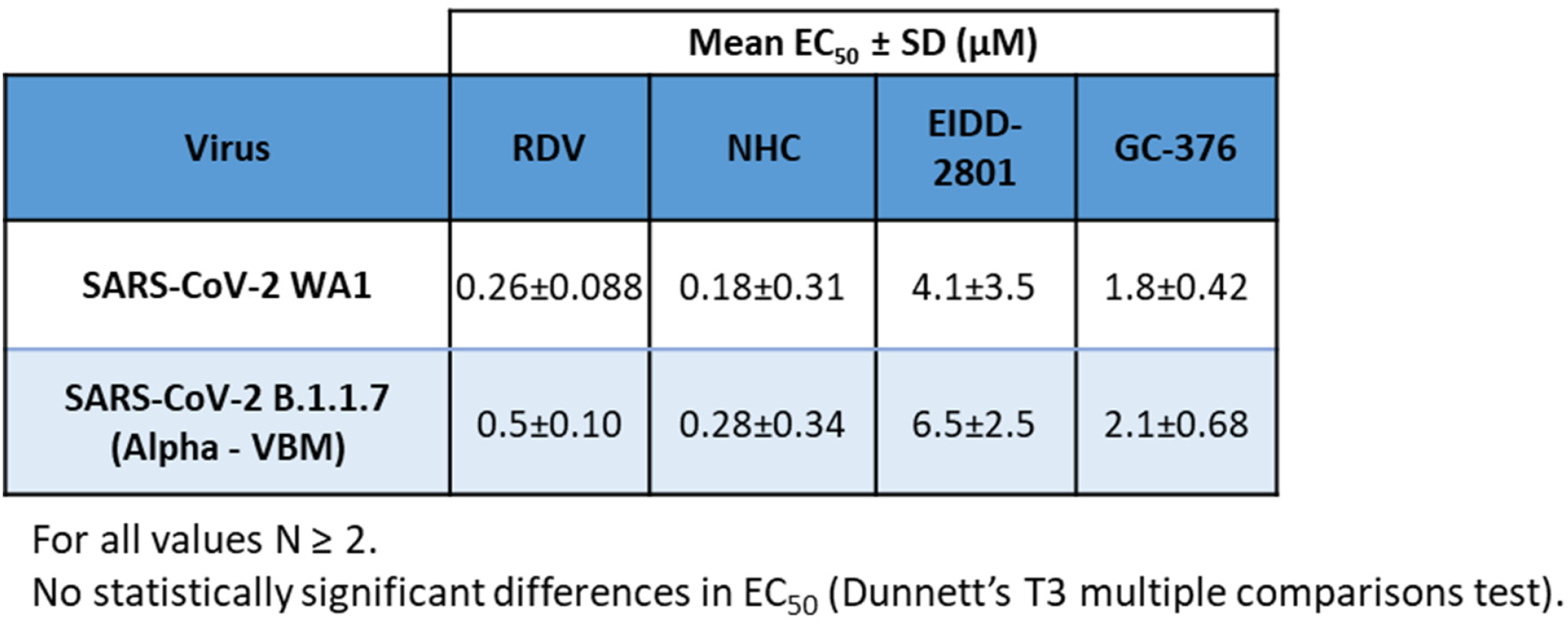
Comparison of B.1.1.7 Variant Being Monitored (VBM) to parental WA isolate on antiviral efficiency in Calu3 cells.

**Figure S1.**
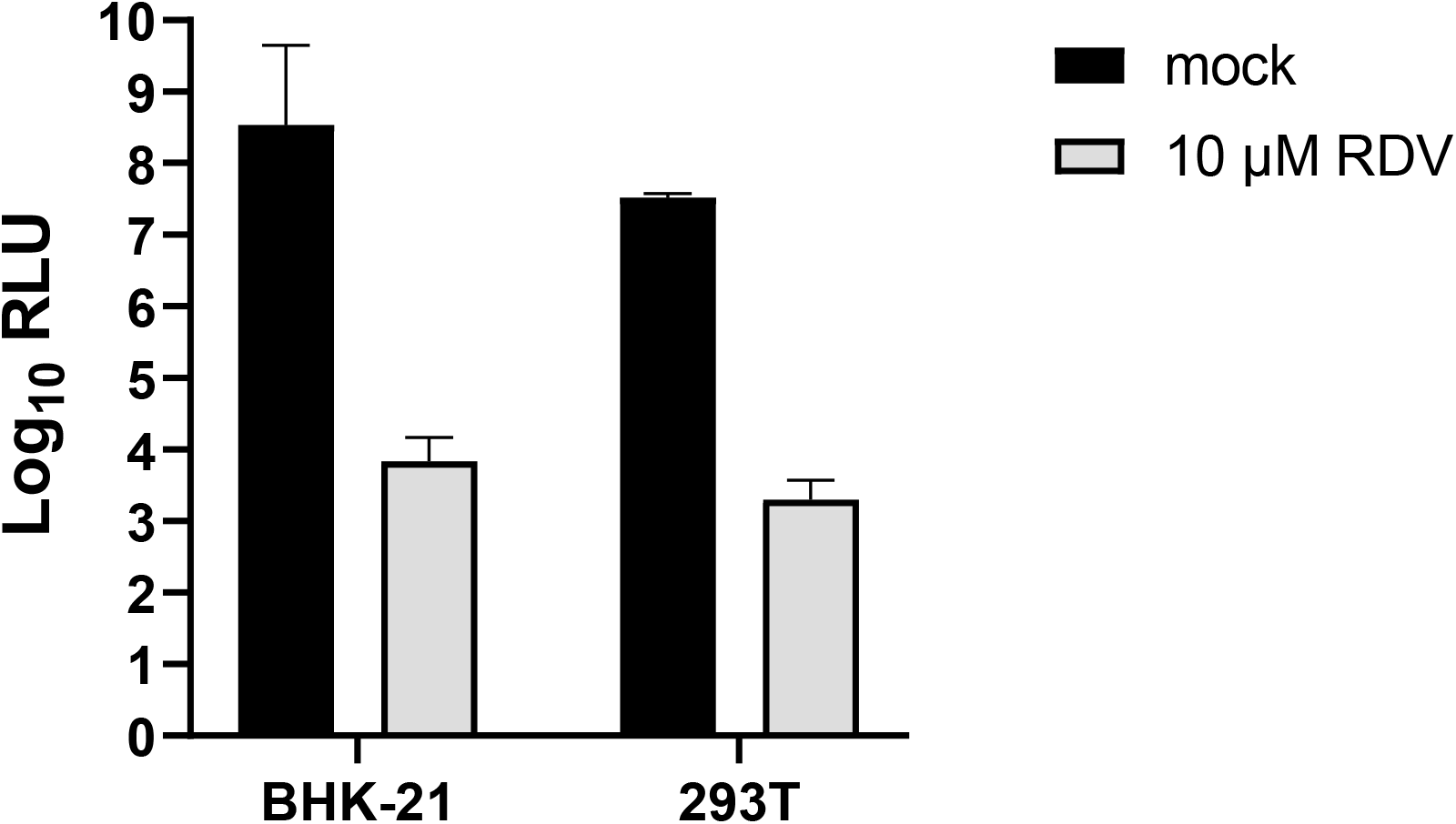
SARS-CoV-2 replicon activity in BHK-21 and 293T cells. SARS-2R_NL_Puro^R^ RNA was transcribed from the BAC using T7 polymerase. BHK-21 or 293T cells were electroporated with SARS-2R_NL_Puro^R^ RNA then seeded into a 24-well plate and half of the samples were treated with RDV. NLuc activity was determined 48 h post treatment. Data are the average of two independent experiments with standard deviation indicated.

**Figure S2.**
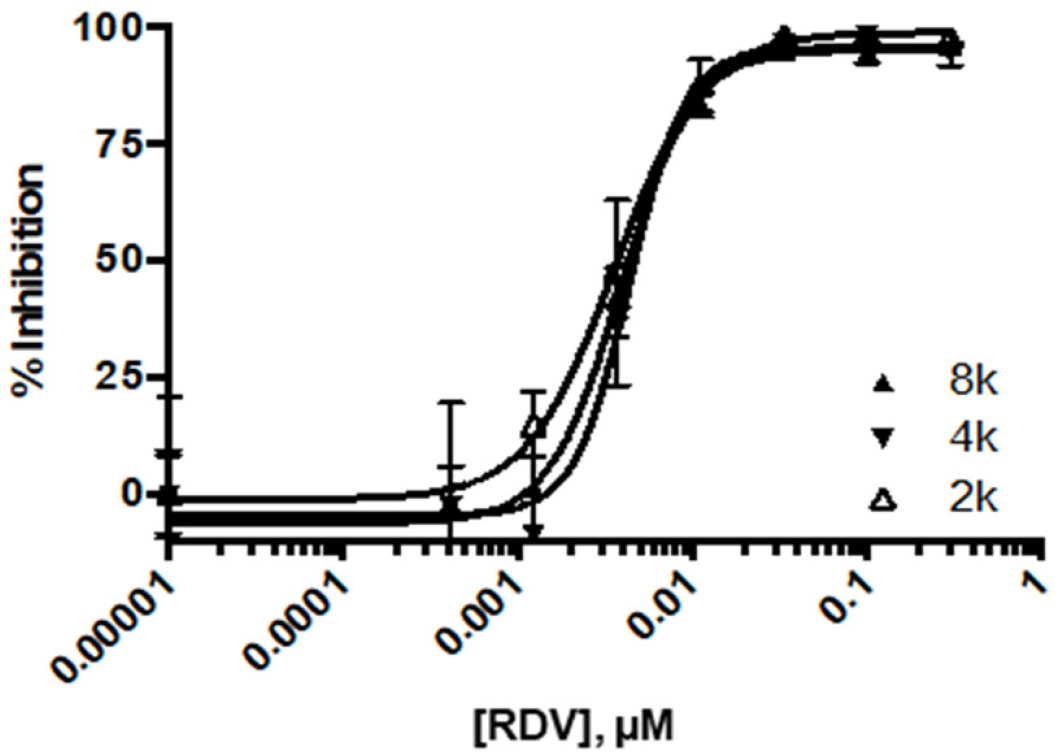
Inhibition of SARS-CoV-2 WT replicon by remdesivir (RDV). 293T cells were transfected with SARS-2R_NL_Puro^R^. Cells were trypsinized then seeded into 384-well plate and treated with RDV. NLuc activity was determined 48 h post treatment and dose response curves plotted.

**Figure S3.**
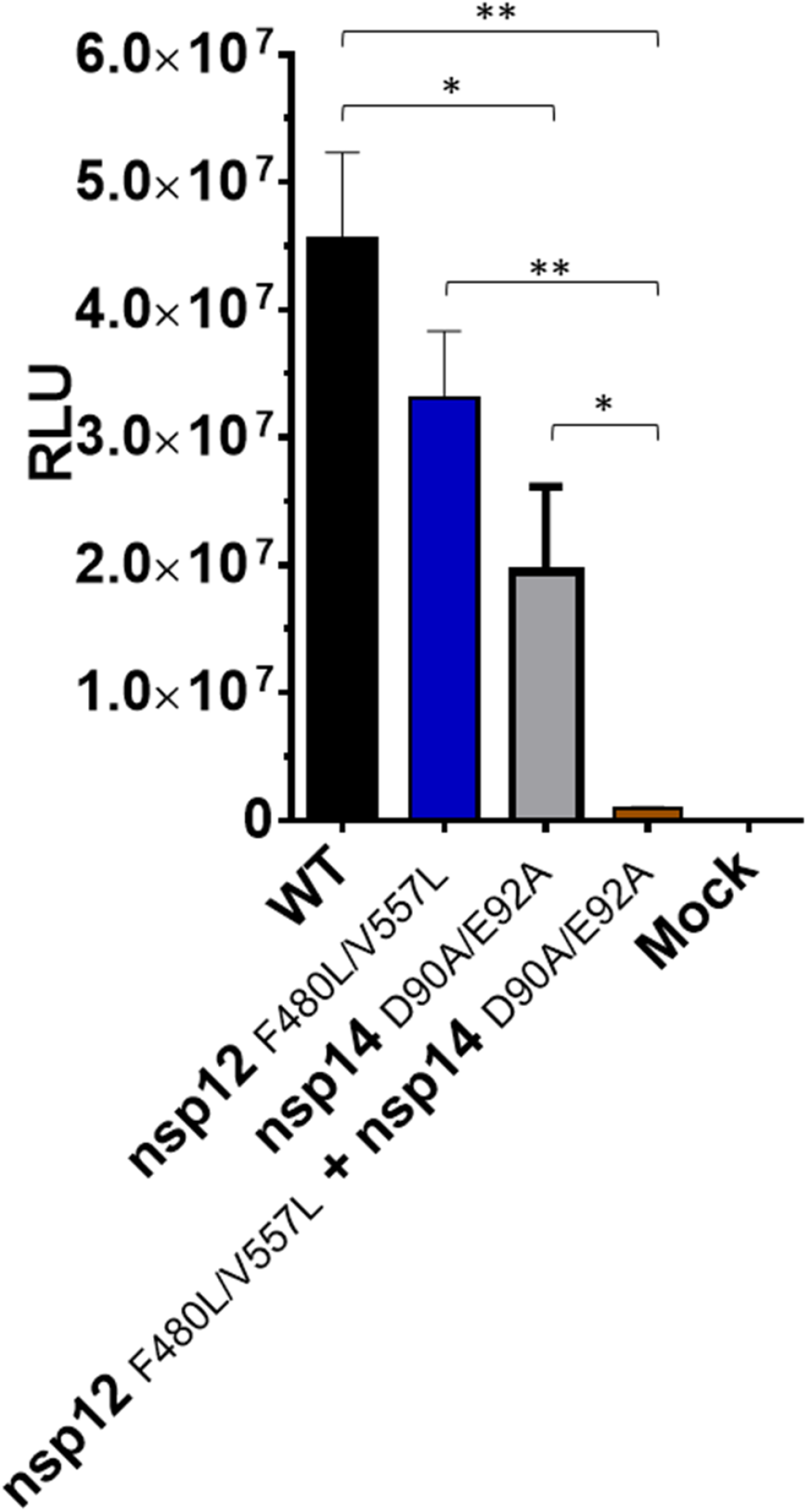
Fitness of SARS-2R-NLuc-Puro^R^ replicons bearing mutations that modulate resistance to RDV. BHK-21 cells were transfected in 24-well plates with 0.25 μg of SARS-2R replicons carrying the indicated mutations, together with 0.025 μg each of N, ORF3b, and ORF6 expression plasmids. NLuc activity was measured at 48 hpt. Averaged data from 2 independent experiments are shown with standard deviations. P values for pairwise comparisons determined by 2-way ANOVA: ** p ≤ 0.01; * p ≤ 0.05.

**Figure S4.**
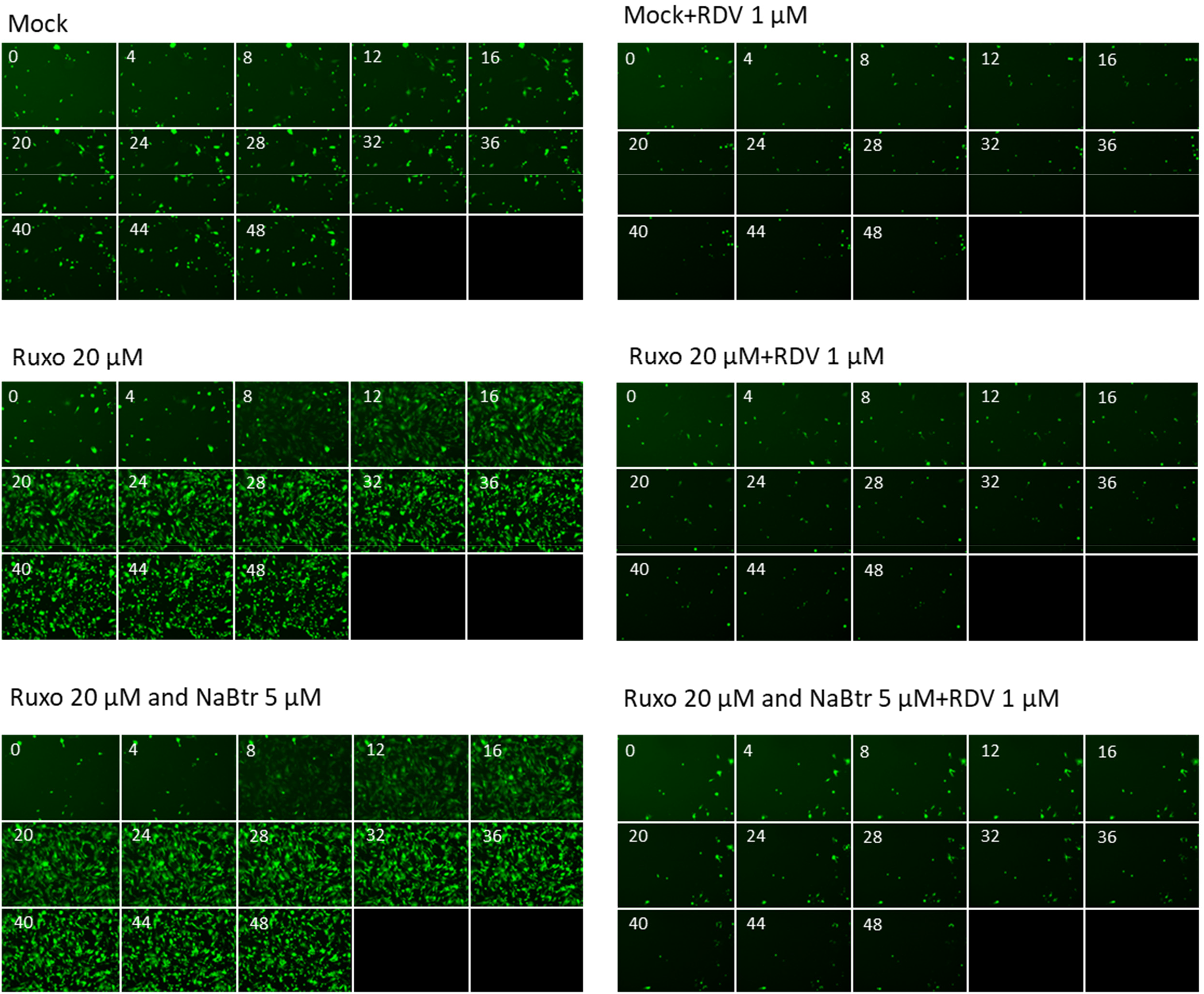
SARS-CoV-2 replicon activity following inhibition of IFN-signalling and derepression of transcription. BHK-SARS-2R_GFP_Neo^R^_NL cells were seeded in a 96-well plate at 10,000 per well. After 24 h, media was replaced with media containing the additives indicated. Replicon activity was assessed by counting GFP-positive cells. Images were collected at 4 h intervals using a high content live-cell imaging system. NaBtr, sodium butyrate; Ruxo, ruxolitinib; RDV, remdesivir. Representative data shown from one of two independent experiments. Data presented here are part of the raw data presented in Figure 5.

**Figure S5.**
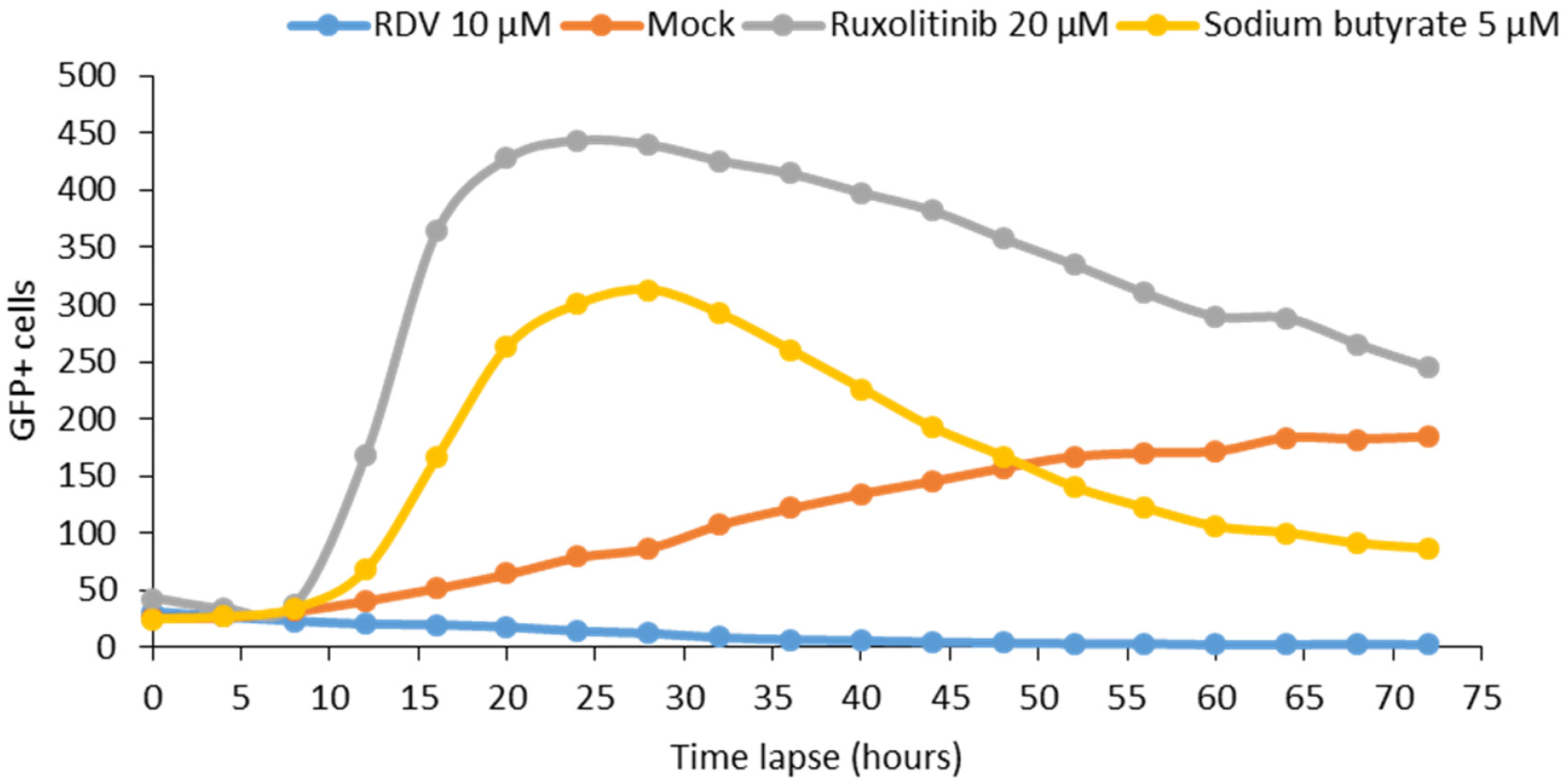
SARS-CoV-2 replicon activity following inhibition of IFN-signalling and derepression of transcription. BHK-SARS-2R_GFP_Neo^R^_NL cells were seeded in a 96-well plate at 10,000 per well. After 24 h, media was replaced with media containing the additives indicated. Replicon activity was assessed by counting GFP-positive cells. Images were collected at 4 h intervals using a high content live-cell imaging system. RDV, remdesivir. Representative data shown from one of two independent experiments.

**Figure S6.**
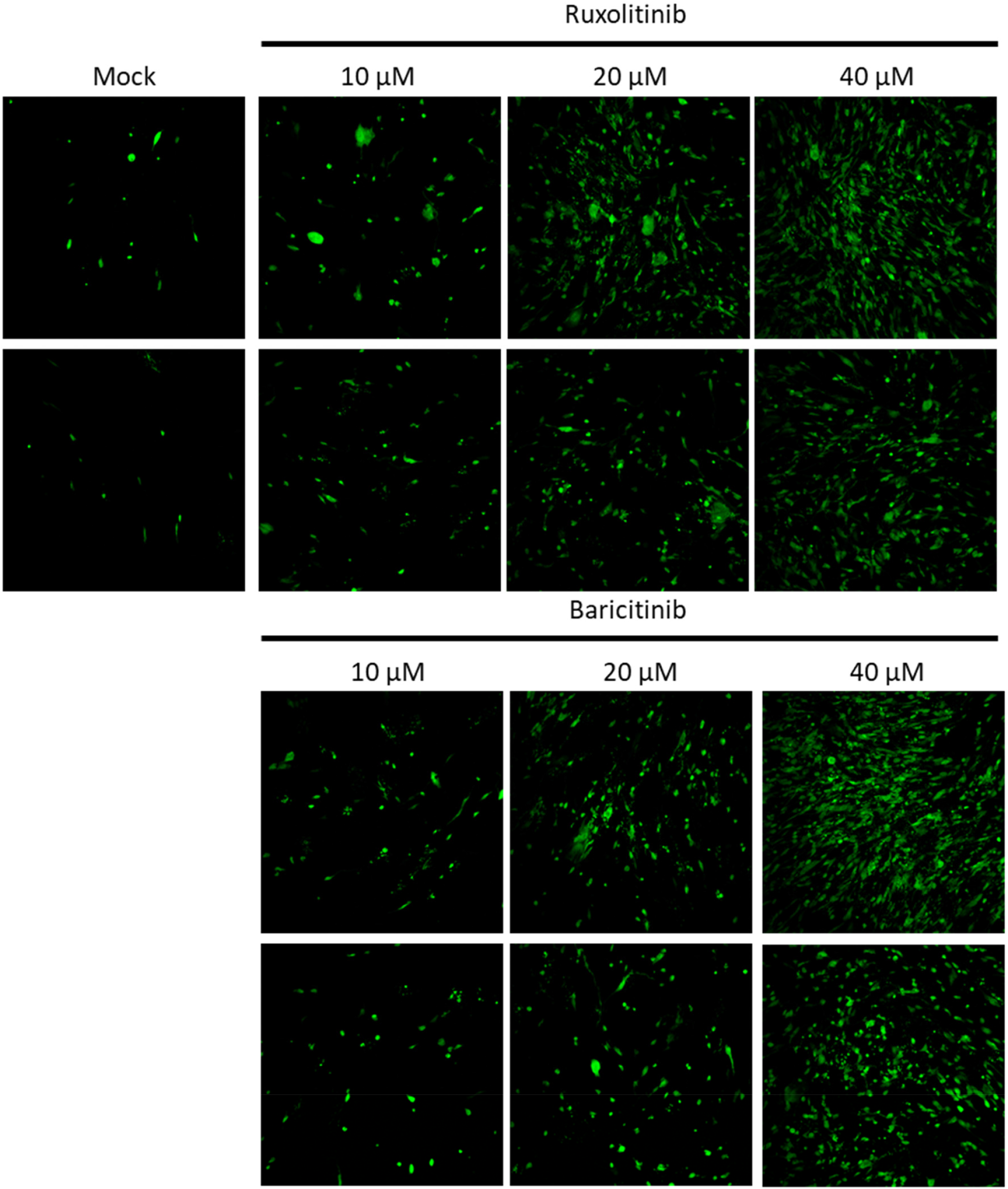
SARS-CoV-2 replicon activity following inhibition of IFN-signalling and derepression of transcription. BHK-SARS-2R_GFP_Neo^R^_NL cells were seeded in a 96-well plate at 10,000 per well. After 24 h, media was replaced with media containing the additives indicated. Replicon activity was assessed by imaging GFP-positive cells after 24 h using a high content live-cell imaging system.

**Figure S7.**
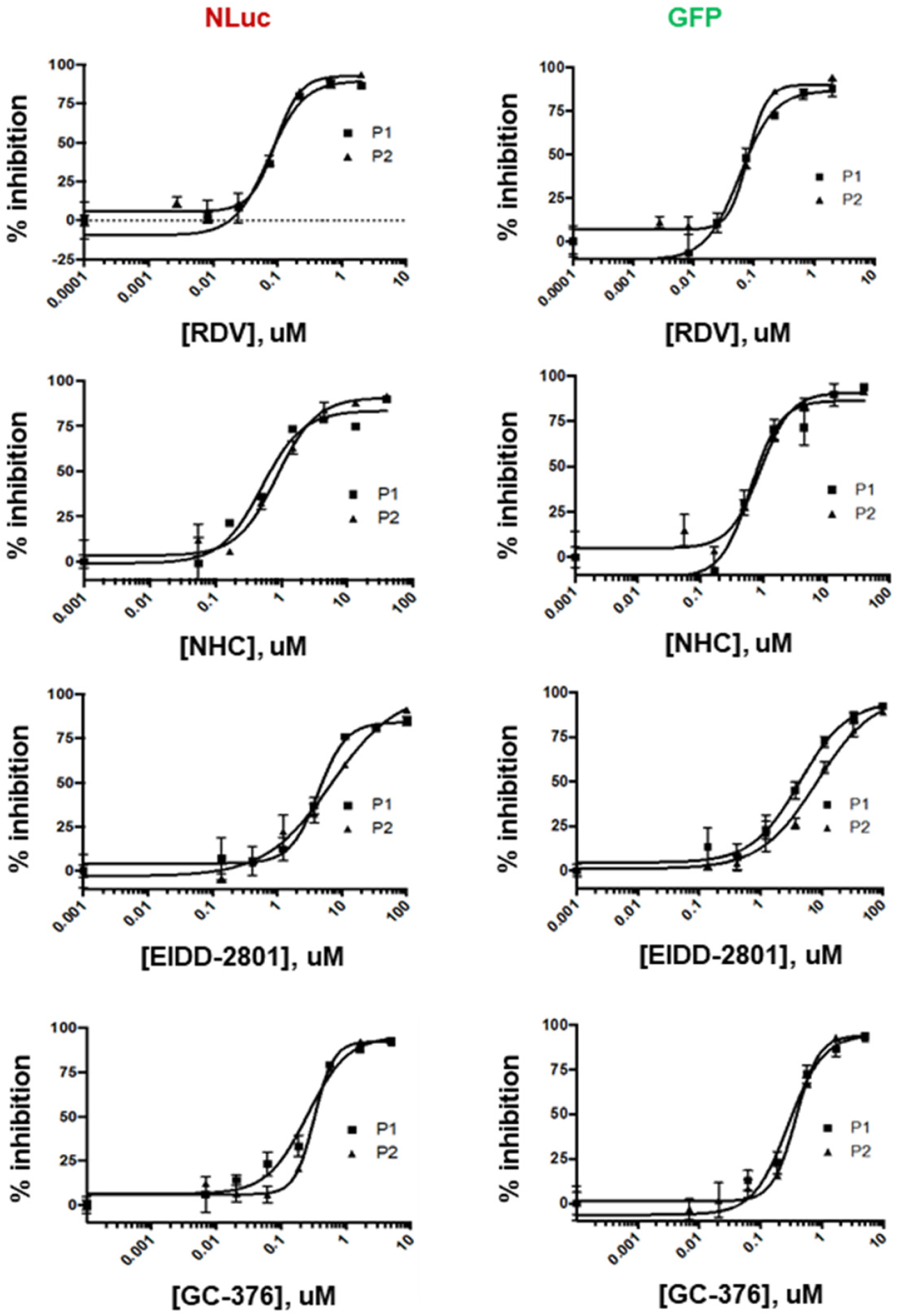
Inhibition of replication activity from BHK-SARS-2R_GFP_Neo^R^_NL replicon-expressing stable cell line as measured by multiplex NLuc and GFP markers. The replicon harboring BHK-SARS-2R_GFP_Neo^R^_NL cell line was challenged with RDV, NHC, EIDD-2801, and GC-376 in 96-well plates. At 48h post treatment, NLuc activity of supernatants from each well were measured and total GFP positive cells were counted. P1, P2: two biological replicate plates.

